# Single-cell and Spatial Omics Reveals Region-Specific Plasticity and Therapeutic Vulnerabilities in Metastatic High-Risk Neuroblastoma

**DOI:** 10.1101/2025.10.19.681936

**Authors:** Lai Man Natalie Wu, Janet L. Oblinger, Dazhuan Xin, Rohit Rao, Feng Zhang, Mike Adam, Oscar Lopez-Nunez, Sara Szabo, Daniel von Allmen, Long-Sheng Chang, Brian D. Weiss, Q. Richard Lu

**Author notes:** Correspondence: Lai Man Natalie Wu and Q. Richard Lu; and.

## Abstract

Neuroblastoma, a deadly pediatric cancer derived from sympathetic ganglia of the peripheral nervous system, frequently metastasizes, driving poor outcomes. Primary neuroblastomas are well-characterized, but the mechanisms underlying metastasis remain poorly understood. Here, by applying single-cell and spatial multi-omics analyses to primary and metastatic tumors, we found that lymph-node metastases in high-risk neuroblastomas display distinctive cellular heterogeneity and plasticity, marked by mesenchymal-like and stem-like states and heightened epithelial-to-mesenchymal transition activity compared to primary adrenal tumors. Additionally, compared to primary adrenal masses, the metastatic niche display increased immunosuppressive myeloid programs, heightened immune checkpoint signaling, and lymphocyte exhaustion, which are indicative of immune evasion and dysfunction. Notably, metastatic neuroblastomas show elevated eIF4F translation machinery and XPO1 levels. Dual inhibition of eIF4A and XPO1 synergistically halted tumor growth and prolonged survival in xenograft models. Together, our multi-omics studies reveal the molecular and cellular plasticity that contributes to therapy resistance and highlight exploitable therapeutic vulnerabilities in high-risk metastatic neuroblastomas.

## INTRODUCTION

Neuroblastomas (NBs), which originate from the sympathetic ganglia derived from the neural crest of the peripheral nervous system, are the most common and most deadly extracranial childhood tumors, accounting for 15% of all pediatric cancer deaths ^1,2^. These tumors arise primarily from the neuroblasts of the neural crest lineage along the sympathetic nervous system with more than half of the cases occurring in the adrenal medulla ^1^. NB is a highly heterogeneous disease with variable clinical behaviors ranging from spontaneous tumor regression to rapid progression, metastasis, and therapy resistance. Patients with low-risk NBs have nearly 100% survival rate with observation and/or resection. Patients with intermediate-risk disease have an overall survival rate of >85% with resection and low-dose chemotherapy, whereas patients with high-risk NBs have <50% overall survival, despite intense multimodal therapy including external beam radiotherapy, immunotherapy, high-dose chemotherapy, and stem cell rescue ^3^. Amplification of the oncogene *MYCN* is the most frequent genetic aberration associated with high-risk NB ^4,5^. A better characterization of tumor heterogeneity and plasticity will improve therapy for NB patients.

Nearly 50% of NB patients have metastatic disease (stage M) at diagnosis, with the bone marrow and bone and regional lymph nodes as the most common sites of metastasis ^6^. Moreover, NB patients with metastatic lesions are at risk for relapse after chemotherapy, and salvage therapies for relapsed patients are often ineffective, leading to dismal patient outcome ^1^. Two core neoplastic states—adrenergic-like (ADRN-like) and mesenchymal-like (MES-like)—have been identified in NB ^7–10^. However, it is unclear what makes NB metastasize and whether NB metastases are distinct from primary tumors arising at the adrenal gland. Multiple single-cell transcriptomics studies of patient NB specimens have focused on establishing the developmental origin of NB by characterizing the malignant cells with reference to cell types and lineage trajectories of the developing adrenal medulla ^11–16^. These studies suggest that primary NB resembles neural crest-derived, committed sympathoadrenal progenitors in the developing sympathetic nervous system ^17^. A population of tumor cells may mimic immature cells of neural crest development with mesenchymal-like features such as Schwann cell precursors ^11–16^. A recent single-cell analysis detected an aggressive transitional tumor cell type with adrenergic and mesenchymal characteristics, that serves as a predictor of poor prognosis, underscoring the phenotypic plasticity within the spectrum of these peripheral neuroblastic tumors ^18^. Recent studies of the tumor microenvironment (TME) landscapes of NBs pre- and post-therapy ^19^ and of bone marrow metastases^13^ revealed the immunosuppressive and dysfunctional features of these TMEs. To date, a comprehensive atlas of the dynamic cancer cell states in high-risk NBs at metastatic lymph nodes remains incompletely understood. Lymph nodes are a key transitional niche —an early site where neuroblastoma cells adapt, engage with immune cells, and acquire traits that promote spread and therapy resistance.

Here, we applied single-cell multi-omics and spatial transcriptomics to primary adrenal NB masses and metastatic lesions such as lymph node metastases. This is the first analysis of the dynamic cancer cell states in metastatic lymph nodes. We characterized site-specific cellular heterogeneity, tumor cell fate transitions, and TME landscapes within the primary tumors and metastatic lymph nodes before and after therapy. Our cohort analysis identified distinct phenotypes of malignant neuroblasts in primary and lymph node lesions, with metastases showing mesenchymal and stemness traits and primary tumors diverging from the chromaffin lineage. Further, we identified unique transcriptional circuits that drive metastases through the epithelial-to-mesenchymal transition (EMT) and immune dysregulation and that are correlated with poor outcomes. Targeting these mechanisms synergistically halted tumor growth and metastasis in mouse and patient-derived xenograft (PDX) models. Taken together, our findings reveal unique metastatic niches in high-risk NBs and identify cell signaling mechanisms correlated with clinical outcome, pointing to potential therapeutic vulnerabilities in treatment-resistant NBs.

## RESULTS

### Cellular landscapes of patient high-risk NBs at primary and metastatic lesions differ

To elucidate the complexity of cellular compositions in NBs from primary and metastatic locations, we analyzed samples from patients with high-risk NB, diagnosed based on *MYCN* amplification (Fig. 1A and Supplementary Table S1), using single-cell multi-omics profiling including single-nucleus RNA-sequencing (snRNA-seq) and single-cell assay for transposase-accessible chromatin using sequencing (scATAC-seq). We analyzed six primary tumor specimens (n=50,572 cells) from adrenal glands and five metastatic masses (n=40,959 cells) from reticuloendothelial lymph nodes using the single-cell Multiome assays on the 10X Genomics platform. These tumor specimens had been surgically resected following standard-of-care induction chemotherapy and were obtained from CCHMC oncology tissue repository. Specimens from the primary tumors and matched lymph node metastases were available for two of these patients. We also included three publicly available treatment-naïve high-risk primary NB single-cell RNA-sequencing (scRNA-seq) datasets (n=3,541 transcriptomes) in our analysis.

**Fig. 1.**
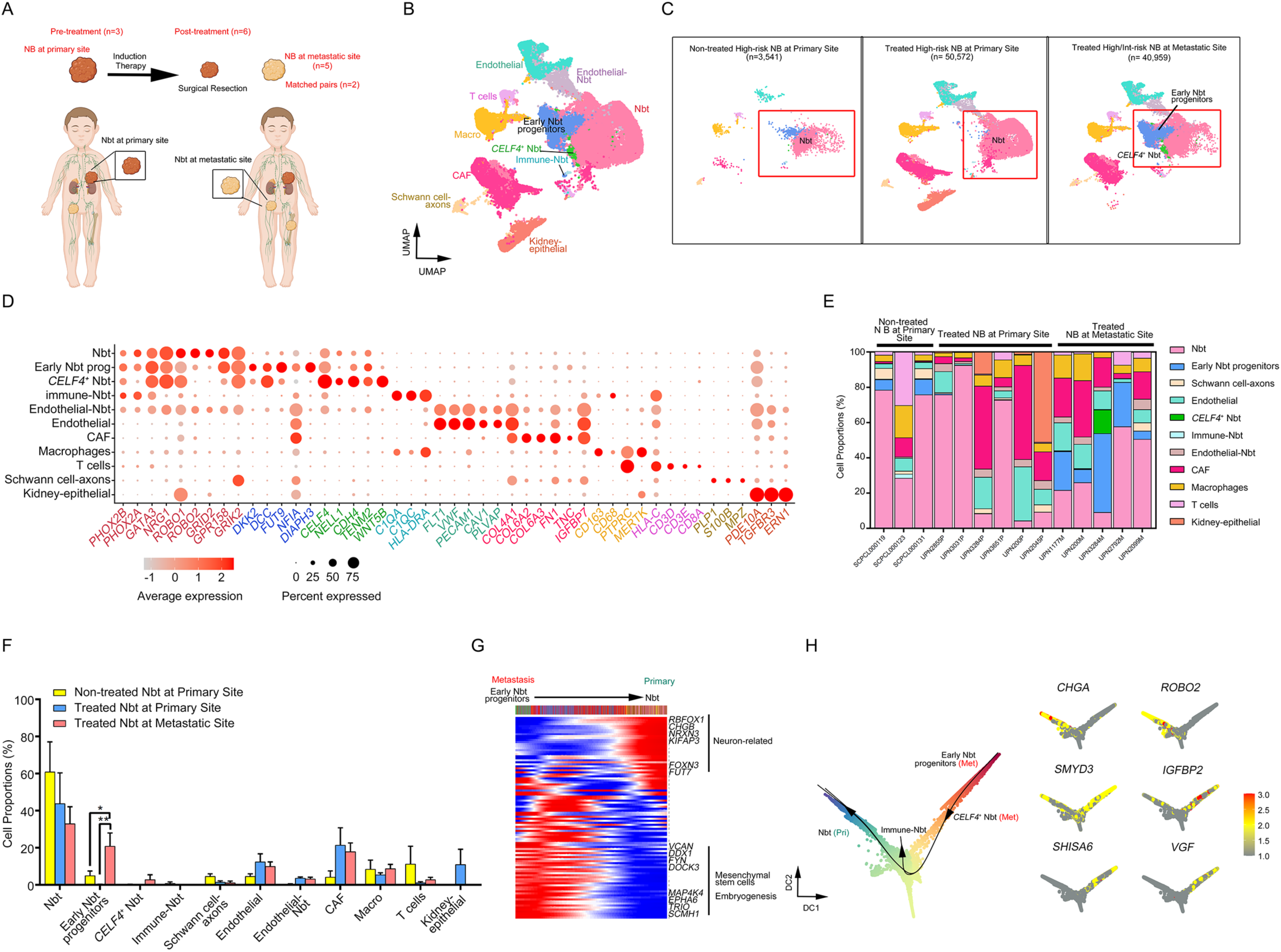
The cellular landscape of high-risk NB at primary adrenal glands and lymph node metastases. a. Schematic overview of included patients and sample collection. scRNA-seq data from pre-treatment biopsies of high-risk NBs (*MYCN* amplified) were obtained from ALSF (Supplementary Table S1). Samples of six NBs resected from primary adrenal glands and five lymph node metastases of patients post-treatment were obtained from CCHMC oncology tissue repository and processed for 10X single-cell multi-omics; both primary and metastatic samples were obtained from two subjects. b. Uniform Manifold Approximation and Projection (UMAP) visualization of single-cell transcriptome data of 95,072 cells from primary tumors from untreated patients, primary tumors resected post therapy, and lymph node metastatic sites resected post therapy. Colors represent assigned cell types. c. UMAP visualizations of single-cell transcriptome data from (left to right) pretreated primary NBs (n=3; 3,541 cells), post-treatment primary NBs (n=6; 50,572 cells), and lymph node metastases (n=5; 40,959 cells). Red boxes indicate tumor cell phenotypes. d. Dot plot of marker gene expression in cell clusters from NB tumors. The color represents scaled average expression of marker genes in each cell type, and the size indicates the proportion of cells that express the marker gene. e. Cell type proportions in primary NBs biopsied pre-treatment and post-treatment primary NBs and metastatic lymph nodes. f. Proportions of cell types in primary NBs biopsied pre-treatment and post-treatment primary NBs and metastatic lymph nodes. \**P* < 0.05, \*\**P* < 0.01; Multiple t-test using the Holm-Sidak method. Data are means ± SEM. g. Pseudotime ordering by FitDevo ^24^ of the developmental potential of NB tumor clusters (from orange as the most undifferentiated state to green as the mature state). h. NB tumor cell state trajectories during tumor progression inferred by Slingshot ^25^. Arrows indicate inverse direction of transformation as tumor cells acquire additional phenotypes at metastasis. i. Heatmap of gene expression dynamics over pseudotime in NB tumor cell state trajectories during transition from primary tumor to metastasis inferred using the GAM function in R ^93^.

To characterize the tumor landscape in localized NBs and lymph node metastases, we integrated single-cell datasets using the Seurat integration platform after quality controls (see methods). We compared the transcriptional profiles of NBs stratified by treatment status and location (Fig. 1B, C and Supplementary Fig. S1). By interrogating the expression patterns of canonical markers for different cell lineages, we cataloged high-quality single nuclei into six major cell types: sympatho-adrenal lineage cells (*PHOX2A*, *PHOX2B*, *GATA3*), Schwann cells (*PLP1*, *S100B*, *MPZ*), stromal fibroblasts (*FN1*, *COL6A2*, *COL4A1*), endothelial cells (*FLT1*, *VWF*, *PECAM1*), myeloid cells (*PTPRC*, *CD68*, *CD163*), and T cells (*CD3D*, *CD3E*, *CD8A*) (Fig. 1D). These key cell lineages were identified in primary and metastatic NBs regardless of treatment status (Fig. 1C-E and Supplementary Fig. S1). Prior to treatment, the primary NB masses were predominantly composed of tumor cells of sympatho-adrenal lineage with minimal presence of immune or stromal components. Following induction chemotherapy, immune and stromal cells became significant constituents of both primary and metastatic NBs, suggesting the development of an inflammatory milieu within tumors after therapy.

Neuroblast-like cells with a strong neuronal signature (i.e., expression of *ROBO1*, *ROBO2*, and *NRG1* ^20^) were the primary components of adrenal masses. In contrast, lymph node metastases had greater tumor cell diversity including novel tumor states enriched with gene signatures associated with cancer stemness and metastasis (e.g., *DKK2*, *FUT9, DIAPH3* ^21–23^) (Fig. 1D-F and Supplementary Fig. S1). The transcriptional profiles of major neuroblast cell states, including neuroblasts and early neuroblast progenitors, also differed between metastatic lesions and primary masses (Supplementary Fig. S2A), suggesting that metastasis results in a considerable shift of tumor cell phenotypes.

To determine the potential tumorigenic trajectories of neuroblast cell phenotypes, we utilized an unsupervised method, FitDevo, which infers developmental potential based on the correlation between sample-specific gene weight and gene expression ^24^. The early neuroblast progenitor cluster, uniquely enriched in neuroblastoma metastases, exhibited a less differentiated phenotype (Fig. 1G-H). We confirmed this finding by the pseudo-temporal ordering of neuroblast cell state evolution via Slingshot using a diffusion map approach ^25,26^. To dissect the expression dynamics along the trajectory, pseudotime analysis was used to order the expression of the most differentially expressed genes between neuroblasts and early neuroblast progenitor populations (Fig. 1I). Several genes that encode proteins involved in cancer invasion and related to EMT, such as *IGFBP2*, *VGF*, and *SMYD3*, were robustly expressed in metastasis-specific clusters, whereas adrenergic-related genes (*CHGA*, *ROBO2*) were more highly expressed in the differentiated cell state within NB tumor cell continuum (Fig. 1I and Supplementary Fig. S2B). This analysis suggests that the acquisition of primitive and less differentiated cell states occurs during metastasis and underscores the importance of lineage plasticity during NB dissemination.

### Cellular heterogeneity increases during NB metastasis

To decipher the dynamics of cellular plasticity in NB metastases, we evaluated neuroblast populations between primary and metastatic NBs post-therapy by unsupervised clustering of transcriptomic data. Diversity of malignant cell states was higher at metastatic sites than in primary tumors (Fig. 2A-B). In contrast to primary masses, which had a robust adrenergic signature and harbored predominantly neuroblast-like clusters, metastatic sites had diverse malignant populations including neural-crest neuroblast (Nbt) progenitors (*TENM2*, *FGF13, DSCAM* ^27^), mesenchymal neural-crest-like tumor cells (MES-Nbt; *EYA1, PTN, PDE3A* ^28,29^), and their cycling counterparts (*TOP2A*, *MKI6, HMGB2)*, as well as a discrete *DKK2^+^* Nbt metastatic phenotype (*DKK2^+^*NB Met; *VCAN, FUT9, DCC, DKK2* ^21,22,23^) (Fig. 2A-E and Supplementary Fig. S3). Thus, there is a high frequency of tumor cells with mesenchymal neural crest identity at the metastatic locations.

**Fig. 2.**
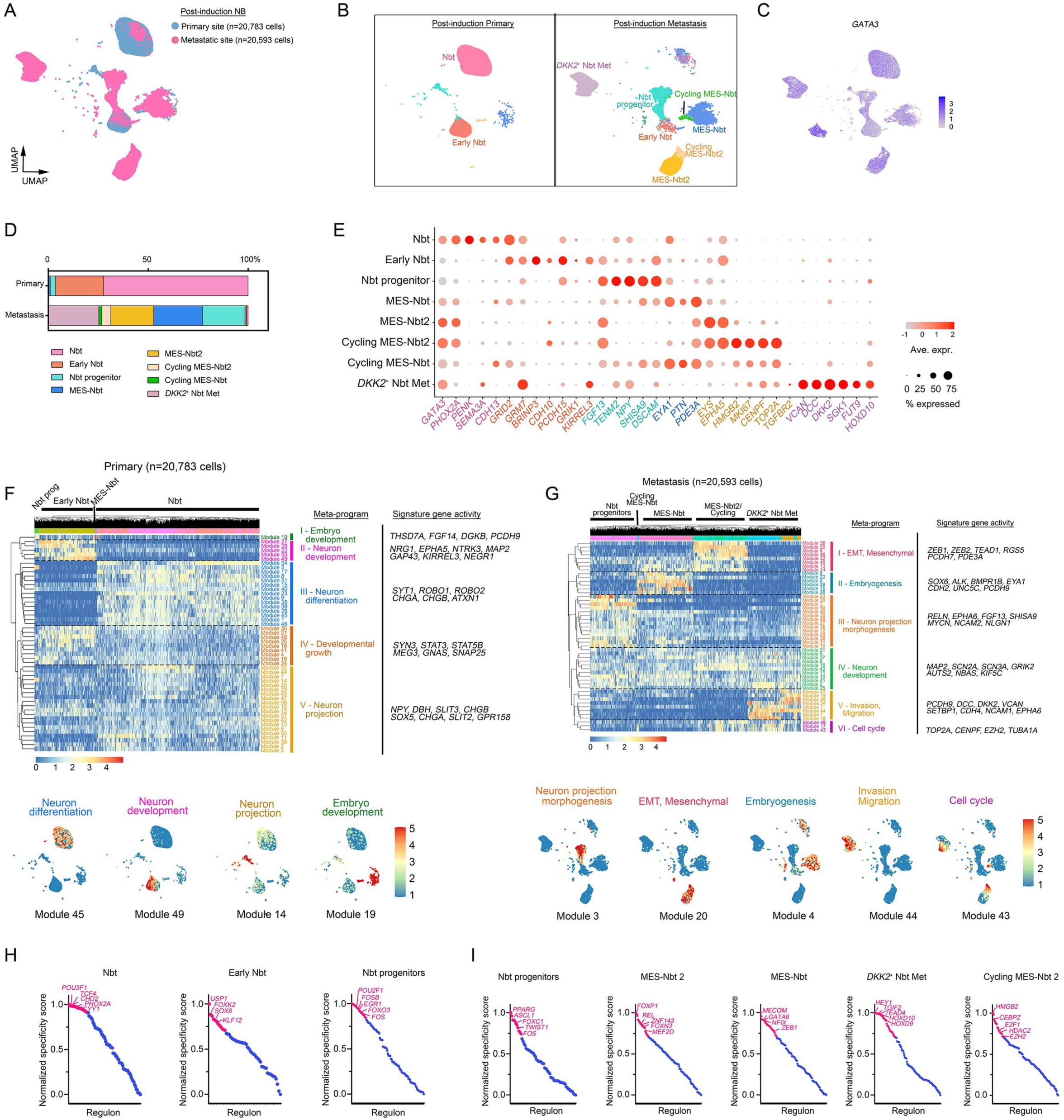
Tumor cell diversity increases upon metastasis. a. UMAP embeddings of NB tumor cell subclusters identified using anchor-based integration by Seurat of transcriptomic data from primary and metastatic tumors, colored by tumor location. b. UMAP visualizations of NB tumor cell subclusters in post-treatment NBs at primary (n=20,783 cells) sites (left) and metastatic (n=20,593 cells) sites (right) after re-clustering. Colors represent assigned cell types within NB tumors. c. UMAP visualization of NB tumor cell subclusters colored by expression of *GATA3*, an NB marker. d. Proportions of identified NB subpopulations at primary and metastatic sites. e. Dot plot of marker gene expression in cell clusters from NB populations. The color represents scaled average expression of marker genes in each cell type, and the size indicates the proportion of cells expressing marker genes. f-g. Upper: Heatmaps of signature expression with cells sorted by non-negative matrix factorization gene modules grouped by key meta-programs in NB cell states at g) primary and h) metastatic sites. X-axis, distribution of NB-derived tumor clusters; y-axis, hierarchical clustering of gene modules into meta-programs. Lower: UMAP visualizations of modules that highlight cell-type specific meta-programs and cellular activity in f) primary and g) metastatic NB populations. h-i. Cell type-specific transcription factor regulons (x-axis) derived by SCENIC plotted against normalized specificity scores (y-axis) in NB cell states at h) primary and i) metastatic sites. Key transcription factors enriched in each cell state are highlighted in magenta.

To further characterize the malignant identity of neuroblast populations, we inferred genome-wide chromosomal copy number variations from the snRNA transcriptomic profiles of primary and metastatic tumor cells using stromal and immune cells as non-malignant cell references ^30^. In tumor cells, we detected genetic alterations frequently reported in high-risk NBs, such as 2p gain ^31^, 6q loss ^32^, 1q and 17q segmental gains, whole-chromosome 7 gains ^33^, and chromosome 3 loss ^34^ (Supplementary Fig. S4), consistent with previous reports of prognostic markers in NBs. These findings further confirm the malignant nature of the tumor populations uniquely identified in NBs at the metastatic sites.

Next, we used CIBERSORTx ^35^ to compare the cells from primary and metastatic sites to reference profiles of sympathoadrenal regions during human embryogenesis from post-conception weeks 6-14 (GEO:GSE147821) ^15^. Deconvolution analyses of cell-type abundances from bulk transcriptomes of human NBs at primary and metastatic sites suggested that primary masses transcriptionally mimicked the sympathoblast signature (Supplementary Fig. S5A), corroborating their sympathoadrenal origin. In contrast, metastatic tumor cell states had less resemblance to sympathoblasts and increased gene signatures representative of immune and mesenchymal lineages (Supplementary Fig. 5A-B).

We next carried out integrated analysis of our snRNA-seq data with reference datasets of human embryonic lineages in the sympathoadrenal regions ^15^. Whereas the stromal and immune lineages in NB samples aligned with embryonic abdominal mesenchymal and immune populations, the tumor cell states distinct from normal embryonic sympathoblasts or chromaffin cells (Supplementary Fig. S5C-F), indicative of NB-specific regulatory aberrations. Furthermore, other than Schwann cell component in the TME, we did not detect Schwann cell precursor-like populations in NB samples.

To characterize the full transcriptional spectrum of neuroblast phenotypic heterogeneity within the NB samples, we applied non-negative matrix factorization (NMF) to define intra-tumor expression programs that consist of genes co-expressed across NB cell states ^36^. We identified five meta-gene activity programs in primary NB tumors and six in metastatic masses that captured dynamic phenotypic alterations consistent with the acquisition of divergent cell identities as these tumors metastasized (Fig. 2F-G). Across primary NB cell states, neuroblasts were highlighted by meta-program III, a sympathoadrenal differentiation signature including expression of *CHGA*, *CHGB*, and *SYT1*. Meta-programs II (*NRG1*, *NTRK3*) and IV (*STAT3*, *GNAS*, *TOP2A*, *CENPF*) represent neuron development and developmental growth signatures, respectively, and were specifically expressed in early neuroblast cells (Fig. 2F). In the tumor cells at metastatic locations, we detected meta-program IV (*MAP2*, *GRIK2*), a hallmark of neuron development, and meta-program I, the mesenchymal-neuroblast-like (MES-Nbt) state, which was enriched for genes related to the EMT such as *ZEB1*, *ZEB2*, and *TEAD1*. Importantly, MES-Nbt cells were also enriched for meta-programs I and II reflective of EMT and embryogenesis, respectively. The *DKK2*^+^ neuroblast-like cells in metastases (Nbt-Met) cells were enriched for meta-program V, enriched for expression of *VCAN*, *CDH4*, and *DKK2*, genes related to invasion and migration. Together, our results suggest that NB tumor cells are plastic and that there mesenchymal-like phenotypes arise during metastasis.

Using SCENIC ^37^, we next reconstructed networks of active transcription factors and their downstream gene targets to identify the gene regulatory networks that govern NB cell state heterogeneity (Fig. 2H-I). We identified key gene regulatory networks relevant to sympathoadrenal lineage specification (e.g., the PHOX2A regulon in neuroblasts and the SOX6 regulon in early neuroblasts), highlighting parallels between normal developmental and NB cell fate determination. In contrast to primary masses, the tumor states in NB metastases exhibited markedly upregulated networks associated with early neural development (e.g., TWIST1 and ASCL1 regulons in neuroblast progenitors), EMT and tumor dissemination (e.g., GATA6 ^38^ and ZEB1 ^39^ regulons in MES-Nbt1 cells and FOXN3 ^40^ in MES-Nbt2 cells), and invasive oncogenic traits (e.g., TGIF2 in the *DKK2*^+^ Nbt-Met state) (Fig. 2I). Thus, tumor location and spread serve as contextual determinants of neuroblast cell states, with committed neuroblasts of the sympathoadrenal lineage predominantly found in primary sites and stem-like neuroblast progenitors and mesenchymal-like tumor cells enriched in lymph node metastases.

### Identification of embryonic and EMT programs as core regulatory circuitries in NB tumor cells at metastatic sites

To define the regulatory mechanisms underlying NB metastasis, we compared the transcriptional regulatory networks of metastatic lesions with those of the primary tumors (Fig. 3A). We performed scATAC-seq and profiled high-quality nuclei from samples from four primary tumors and four lymph node metastases to analyze chromatin accessibility and regulatory dynamics. The single-nucleus data met the expected quality of standard single-cell studies (Supplementary Fig. S6A-B). By integrating snRNA-seq and scATAC-seq datasets of the same nuclei preparations using the ArchR package with batch effect correction ^41^, we annotated all major cell types based on their epigenomic profiles that corresponded to their unique gene expression profiles (Fig. 3A-B and Supplementary Fig. S6C). Based on corresponding gene expression profiles, neoplastic cells were categorized into distinct epigenomic states including adrenergic (i.e., Nbt and early Nbt states), neural crest, and mesenchymal (i.e., Nbt progenitors and MES-Nbt) (Fig. 3B-D). These data indicate substantial differences in chromatin landscapes in diverse neoplastic populations in NB metastatic lesions. In the metastasis-associated neoplastic cell states, we detected robust expression of neural crest (*ZIC1*, *OLIG3*), mesenchymal (*MSX2, GDF6*), and metastasis-related (*DKK2*) genes in addition to adrenergic genes (e.g., *PHOX2B*) that were shared with primary tumor cell states (Fig. 3C). By peak-to-gene linkage, which leverages integrated scATAC-seq and scRNA-seq for peak accessibility and gene expression correlation, we identified differentially accessible chromatin regions and putative enhancer-to-promoter interactions enriched in gene loci encoding genes associated with embryonic morphogenesis (*NBAS*, *ALK*, *HOXD3* and *HOXD9*), neural crest development (*ZIC2*, *PAX6*), and mesenchymal traits (*GDF6*, *PDGFA*, *SOX9*) in metastasis-specific neoplastic cell states (Fig. 3C-E and Supplementary Fig. S6D).

**Fig. 3.**
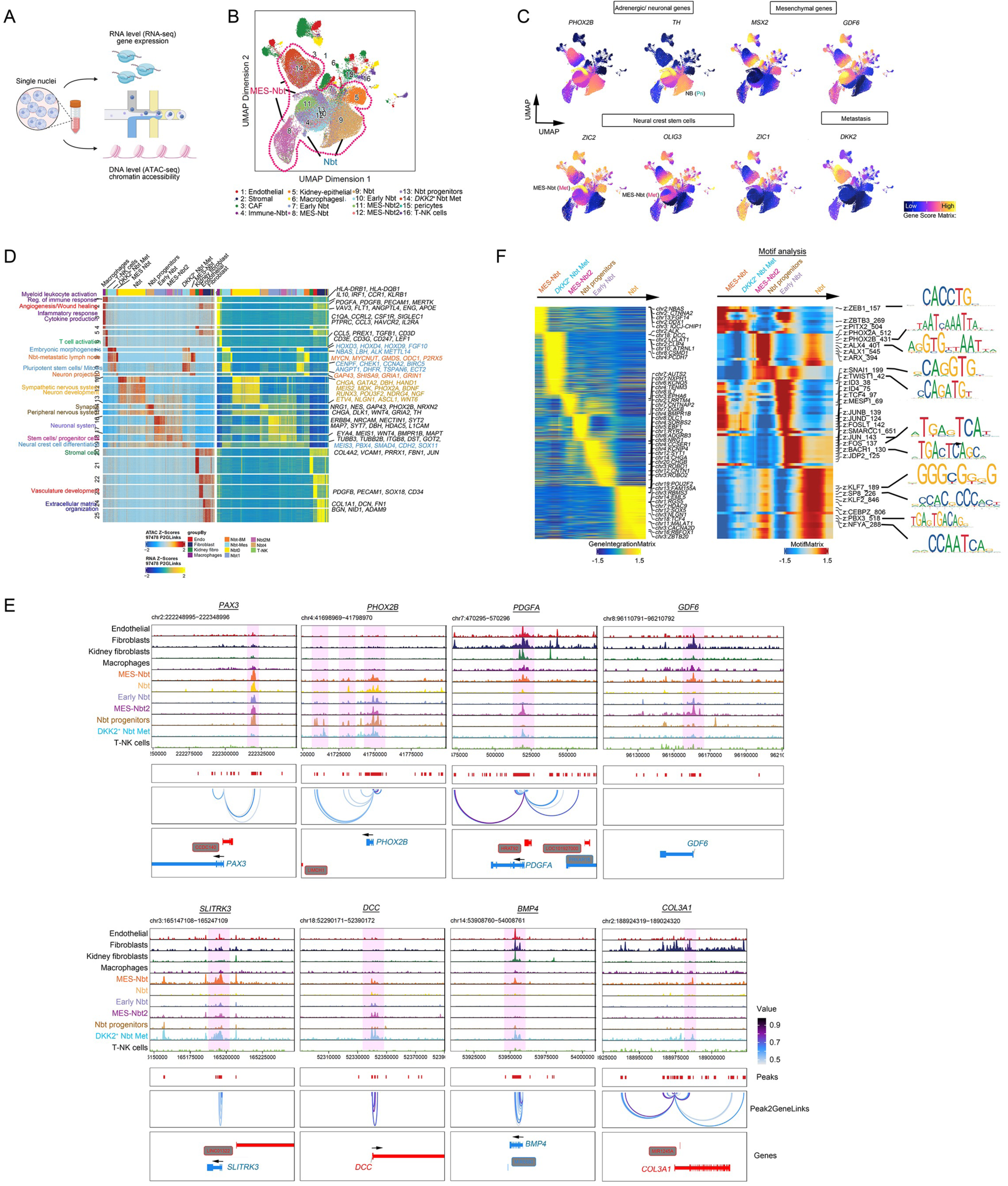
Regulatory networks of stemness and EMT are enriched in tumor cell states in NB metastases. a. A workflow of integrative scRNA and scATAC-seq analysis of gene regulatory networks driving tumor cell states in high-risk NBs. b. UMAP plot of snATAC-seq-derived clusters in NB from both primary and metastatic sites (n=4 samples for each). c. UMAP visualizations of gene accessibility scores of adrenergic, mesenchymal, stem cell, and metastasis markers in distinct NB subclusters. d. Heatmaps of peak-to-gene links identified across cell clusters in NB datasets at primary and metastatic locations. e. Genome browser tracks of co-accessibilities at marker gene loci in NB-derived cell states. f. Heatmaps of differentially expressed gene activity scores (left) and over-represented transcriptional factor motifs (right) in NB tumor clusters in NBs at primary and metastatic sites. Selected motif consensus sequences are shown.

To characterize the gene regulatory mechanisms underlying the emergence of epigenomic states, we defined gene and transcription factor motif score differences between neoplastic cell states present in primary and metastatic sites. In the neuroblast cluster from primary masses, consistent with committed sympatho-adrenergic character, genes with accessible peaks were associated with neuronal differentiation and had higher motif accessibility for adrenergic transcription factors such as PHOX2A and PHOX2B (Fig. 3F). In metastasis-enriched neoplastic populations, such as neuroblast-like progenitors and the MES-Nbt population, genes with open chromatin were those related to activated neural crest specification and EMT programs as indicated by ZEB1, TWIST1, and SNAI1 motif accessibility.

Differential gene expression and gene set enrichment analysis (GSEA) of bulk transcriptomes from primary and metastatic locations revealed a marked upregulation of embryonic stem cell, EMT and mesenchymal signatures, and BMP and TGFβ signaling pathways in tumor cells at metastatic sites (Fig. 4A-D), which contribute to tumor malignancy and invasiveness in NB metastases. Additionally, gene ontology (GO) terms associated with metabolic pathways, such as hypoxia, were markedly enriched in metastases. Conversely, primary NB tumor cells had strong neuronal identity with enrichment in neuronal differentiation gene expression (Fig. 4E). These changes suggest that a shift towards MES phenotype during NB metastasis, which aligns with an acquisition of mesenchymal and embryonic gene regulatory circuitries, resulting in cancer cell reprogramming as NBs metastasize to the lymph nodes.

**Fig. 4.**
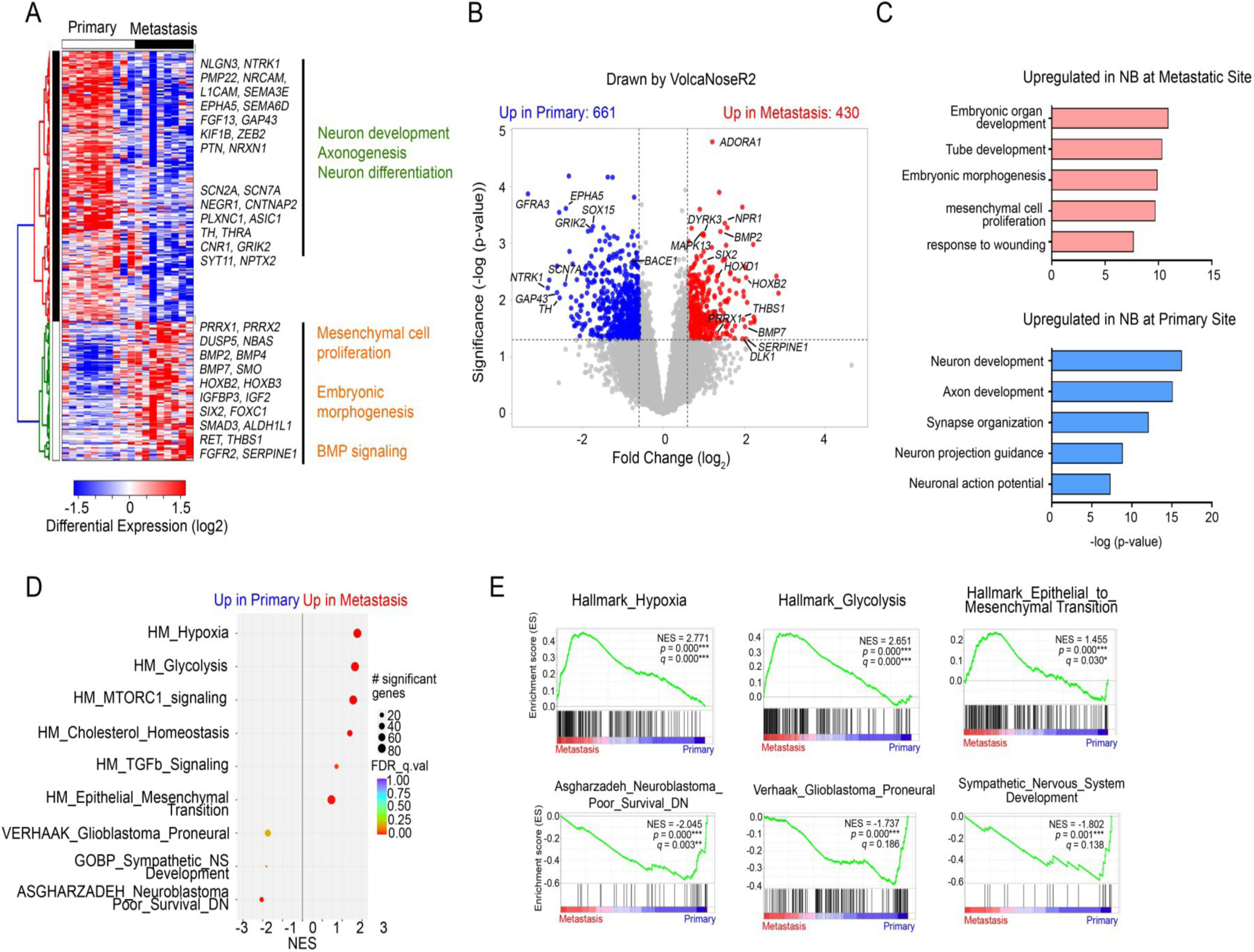
Embryonic and mesenchymal developmental signatures are enriched at metastatic sites. a. Heatmap of differentially expressed genes in NB metastases (n=10) compared with primary tumors (n=8) based on analyses of bulk RNA-seq data. b. Volcano plot of transcriptome profiles of primary and metastatic sites. Red and blue dots represent genes significantly upregulated in metastatic and in primary NB samples, respectively (adjusted *P* < 0.1). c. Top upregulated GO terms in NBs at metastatic (pink) and primary (blue) sites. d. Top differentially regulated gene sets in primary and metastatic NBs based on GSEA analyses. e. GSEA plots of gene signatures upregulated in NB metastases (top panels) and primary tumors (bottom panels).

### Metastatic NBs harbor a higher proportion of immunosuppressive myeloid cells

We next compared immune populations between primary and metastatic locations by analyses of the myeloid compartments of transcriptomic datasets of primary NBs (14 samples) and lymph node metastases (5 samples) resected post treatment (Supplementary Table S1). Subclustering identified 12 functional myeloid-derived states with distinct gene signatures (Fig. 5A-B). These states include classical monocytes (*AQP9*, *EREG*, *SLC2A3*), stress-response macrophages (*HSPA1A*, *HSPA1B*, *APOE*), M2 macrophages (*CD163*, *CD163L1*, *MRC1*), cDC1 (*IDO1*, *CLEC9A*, *FLT3*), and neutrophils (*S100A8*, *S100A9*, *FCN1*) (Fig. 5A-D). Notably, the lymph node metastases harbored a higher proportion of pro-tumorigenic M2 macrophages than did the primary tumors, which were enriched with stress-response macrophages. The gene module scores of both M1 and M2 macrophage signatures ^42^ across all myeloid states and M2 marker staining confirmed strong enrichments of these pro-tumorigenic features in the metastatic lesions relative to the primary tumors (Fig. 5E-F).

**Fig. 5.**
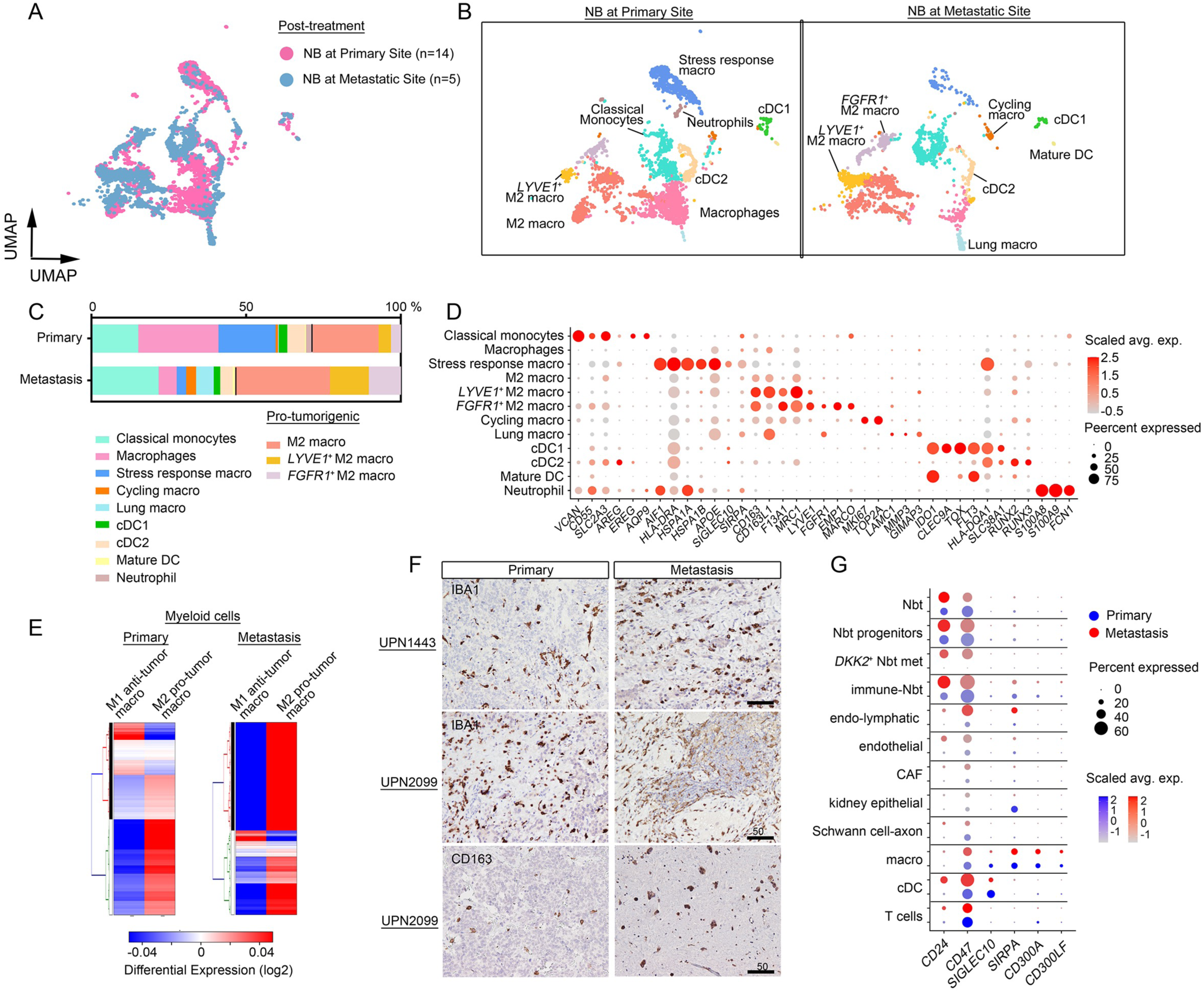
NB tumors are populated with immunosuppressive macrophages. a. UMAP embeddings of myeloid phenotypes identified by anchor-based integration by Seurat of transcriptomic data from post-treatment samples from primary (n=14) and metastatic (n=5) tumors, colored by tumor location. b. UMAP visualizations of myeloid populations from primary (left) and metastatic (right) sites. c. Proportions of myeloid populations in primary and metastatic NBs. d. Marker gene expression in myeloid populations in NBs from primary and metastatic tumors. The color represents scaled average expression of marker genes in each cell type, and the size indicates the proportion of cells expressing marker genes. e. Heatmaps of gene module scores of M1 and M2 macrophage signatures across all myeloid cells from primary and metastatic NBs. f. Representative IHC images of pan- and M2 macrophage markers, IBA1 and CD163, respectively, in primary and metastatic tumors from the same patients. Scale bar, 50 µm. g. Expression of genes encoding ‘don’t-eat-me’ signals in primary and metastatic NBs.

There was more robust expression of the *CD47* and *CD24,* which encode immune checkpoint molecules also known as ‘don’t eat me’ signals ^43,44^, and of genes encoding their corresponding receptors, *SIRPA* and *SIGLEC10*, in macrophages in metastases compared to primary tumors (Fig. 5G). These observations suggest that metastatic tumors exhibit an enhanced evasion of macrophage phagocytosis via expression of ‘don’t eat me’ signals, which convey an inhibitory response in macrophages^43,44^.

### Upregulation of immune exhaustion programs in NBs at lymph node metastases

Lymphocyte dysfunction plays a critical role in fueling cancer growth and progression ^45^, therefore we also evaluated the lymphocyte compartments from the datasets of primary NB and lymph node metastases. We identified seven distinct lymphocyte-derived states (Fig. 6A-C). Lymph node metastases constituted a higher proportion of tumor-associated lymphocytes (2.7%) than primary masses (1.0%) (Fig. 1D-E), as confirmed by immunostaining of CD3^+^ T cells (Fig. 6D). However, lymph node metastases harbored reduced proportions of activated CD8^+^ T cells (*CD8A*, *GZMA*, *CD69* ^46^) and natural killer (NK) cells (*NKG7*, *GNLY*, *PRF1*) (Supplementary Fig. S7A) compared to primary NB masses. By gene module expression analysis, both lymphocytes and NK cells from metastases had diminished activation compared to those from primary tumors (Fig. 6E-F). In addition, NK cells in NB metastases had an enhanced exhaustion signature and prevalent expression of exhaustion regulators such as *TOX* ^47^, *TOX2* ^48^, and *NFAT5* ^49^ when compared with those in primary masses (Fig. 6G-H). Moreover, in NB metastases, lymphocyte exhaustion drivers *TGFBR1* and *TGFBR2* ^50^ were elevated across multiple lymphocyte phenotypes (Fig. 6I and Supplementary Fig S7B). These observations are indicative of heightened immune dysfunction and lymphocyte exhaustion in NB metastases.

**Fig. 6.**
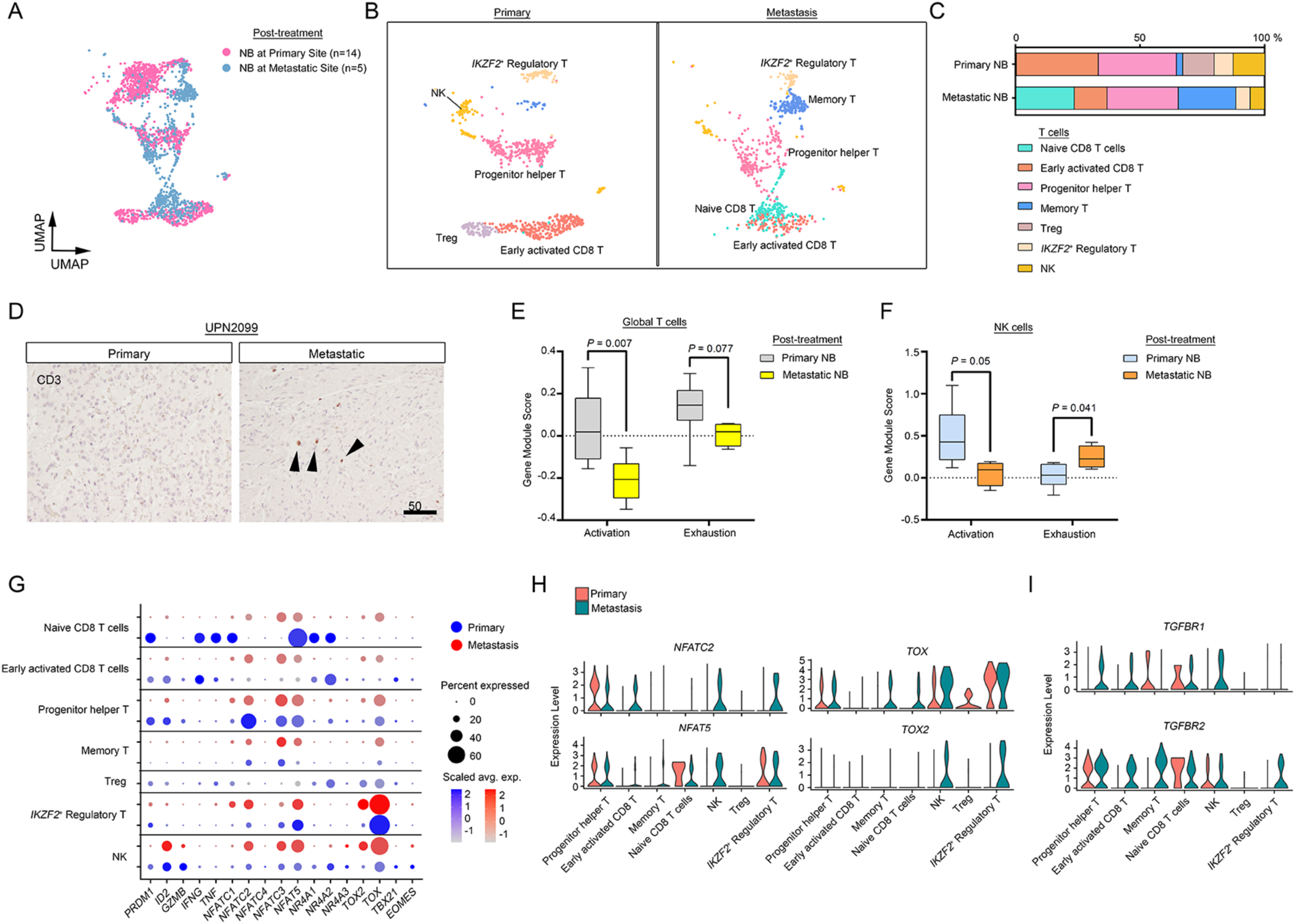
Immune exhaustion programs are enriched in NB metastases. a. UMAP embeddings of T and NK populations identified by anchor-based integration by Seurat of transcriptomic data from post-treatment samples from primary (n=14) and metastatic (n=5) tumors (reference), colored by tumor location. b. UMAP visualizations of T and NK populations from primary (left) and metastatic (right) sites. c. Relative proportions of T cell and NK cell subclusters in primary and metastatic NBs. d. Representative IHC images of pan-T cell marker, CD3, in sections from the same patient. Scale bar, 50 µm. e-f. Gene module scores of lymphocyte activation and exhaustion signatures in e) T and f) NK cells from primary and metastatic sites. g. Percent of cells at primary and metastatic sites that express indicated lymphocyte transcription regulators. h. Violin plots of T cell exhaustion transcription regulator expression in primary and metastatic NBs. i. Violin plots of *TGFBR1* and *TGFBR2* expression in primary and metastatic NBs.

### Spatial transcriptomics analyses identify targetable paracrine signals in NB metastases

Single-cell transcriptomics after tissue dissociation destroys information regarding spatial context that is necessary for understanding of the TME architecture ^51^. To investigate the TMEs in NB metastases compared to primary tumors, performed spatial transcriptomic profiling on a paired primary tumor and matched lymph node metastasis from our cohort using the Visium CytAssist technology from 10X Genomics. Deconvolution analysis of spatial transcriptomics data based on our snRNA-seq-derived signatures identified tumor, immune, and stromal compartments. In the primary NB specimen, we identified two major tumor populations, neuroblasts and differentiating neuroblast-like cells. Both populations expressed classical adrenergic markers *NPY* and *DBH* in the core of the tumor, and this core was surrounded by macrophages, cancer-associated fibroblasts, and Schwann cell-lineage components in the periphery (Fig. 7A-D). Consistent with snRNA-seq data, spatial mapping showed that T cells were scarce within the tumor (Fig. 7D).

**Fig. 7.**
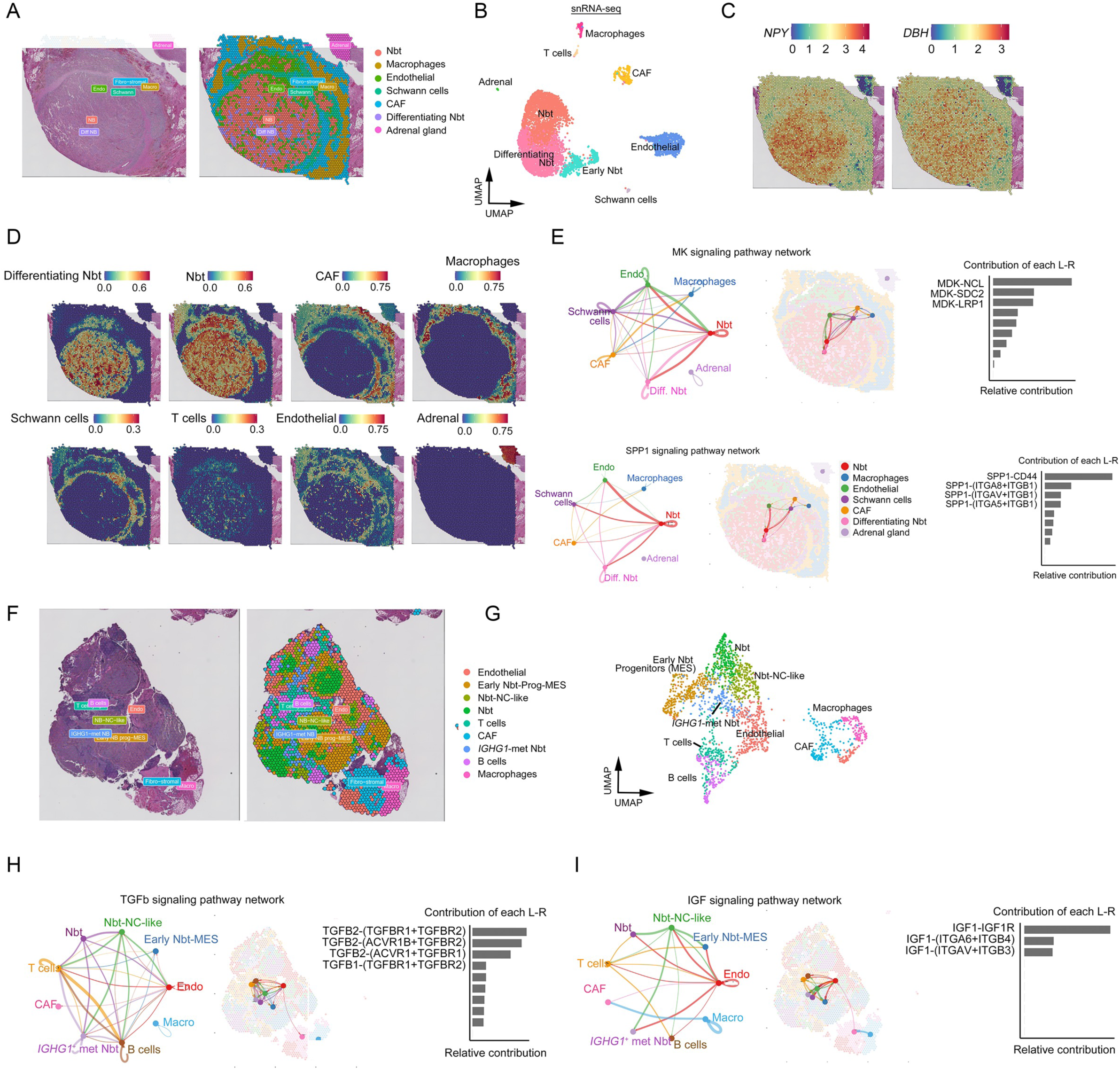
Spatial transcriptomic landscape of primary NBs and metastases identifies potentially targetable paracrine signals. a. Image of a representative H&E section from a primary tumor (left; UPN-2855P) and corresponding spatially resolved transcriptomic data (right) colored by clusters identified using Seurat. b. UMAP visualization of cellular clusters based on snRNA-seq data of 6,119 cells from the primary tumor. c. Primary tumor section colored based on spatial transcriptomics analyses of *NPY* (left) and *DBH* (right). d. Predictions of localization of indicated cell types in primary NB tumor section from anchor-based integration of snRNA-seq data. e. Analyses of inferred intercellular communication networks of MK (top) and SPP1 (bottom) signaling pathways in primary NB. Networks (left) and spatial plots (middle) diagram cell types involved and locations within the tumor section, respectively, and bar charts (right) show the relative contributions of each ligand-receptor (L-R) pair to the communication network. f. Image of representative H&E section from metastatic site (left; UPN-1177M) and corresponding spatially resolved transcriptomic data (right) colored by clusters. g. UMAP representation of nine color-coded clusters defining regional transcriptome diversity in the metastatic site. h-i. Analyses of inferred intercellular communication networks of h) TGFβ and i) IGF (i) signaling pathways in metastatic NB. Networks (left) and spatial plots (middle) diagram cell types involved and locations within the tumor section, respectively, and bar charts (right) show the relative contributions of each L-R pair to the communication network.

The TME is critical for NB growth and metastasis ^52^. To map the TME networks, we integrated snRNA-seq and spatial transcriptomics data and utilized CellChat ^53^ to infer spatially proximal cell-cell communications based on potential for ligand-receptor interactions. We identified a diverse cell-type-specific signaling network composed of secreted ligands and their receptors (Supplementary Fig. S8). These pathways were active at different levels and were mediated by different cell types in primary masses and in lymph node metastases. In primary NB masses, tumor cells predominantly released midkine (MK), a heparin-binding growth factor that enhances tumor cell growth, survival, metastasis, and angiogenesis ^54,55^, to neighboring neuroblasts, Schwann cells, and endothelial cells (Fig. 7E). In addition, tumor cells secreted SPP1 (often called osteopontin), which regulates tumor progression and correlates with poor prognosis in various cancers^56^. This suggests that NB tumor cells produce growth-promoting cues that fuel angiogenesis, progression, and malignancy.

The lymph node metastasis had more heterogeneous tumor cell states and the tumor cells were in closer proximity to T cells and B cells than in the primary tumor sample (Fig. 7F-G). A unique tumor cluster was identified in the metastatic sample that strongly expressed metastasis-associated marker *IGHG1*, which encodes a factor that promotes oncogenesis and progression ^57^. In agreement with robust *TGFBR1* and *TGFBR2* expression in lymphocytes of the metastatic lesions, our spatial ligand-receptor analysis revealed active immunosuppressive TGFβ signaling ^50^ mediated by tumor cells targeted lymphocyte populations (Fig. 7H). This result suggests that TGFβ signaling mediates immune exhaustion in metastatic NBs. IGF-mediated signaling was also enriched in metastatic NB (Fig. 7I); this pathway is critical for EMT and tumor growth ^58^. Collectively, our analyses indicate that NB lymph node metastases possess distinct TME characteristics and cell-cell communication networks and exhibit unique intra- and inter-cellular growth signaling niches compared to primary NBs.

### Dual inhibition of protein translation initiation and nuclear export is a potential strategy for NB treatment

*MYCN* amplification is the primary oncogenic driver associated with metastatic disease and dismal survival in high-risk NB ^1^. Amplification of *MYCN* results in deregulation of the protein translation machinery by enhancing protein synthesis via the eIF4F translation initiation complex, which is composed of eIF4E, eIF4G, and eIF4A ^59^. Elevated levels of eIF4F components are correlated with higher tumor grade and with metastasis ^60^. In snRNA-seq datasets, we detected higher levels of expression of genes encoding the eIF4F components in metastatic NB masses compared to primary tumors (Fig. 8A-D). Consistent with this, spatial transcriptomic profiling revealed stronger expression of *eIF4A1* and *eIF4G1* in the metastatic sample than in the primary tumor sample (Fig. 8E-F). We confirmed widespread cytoplasmic and nuclear localization of eIF4F complex components in NB tumor sections by immunohistochemistry (Supplementary Fig. S9A). Interestingly, the elevated levels of *eIF4A1, eIF4E1*, and *eIF4G1* were associated with poor prognosis of NB patient cohorts^61^ (GSE45547; Fig. 8G).

**Fig. 8.**
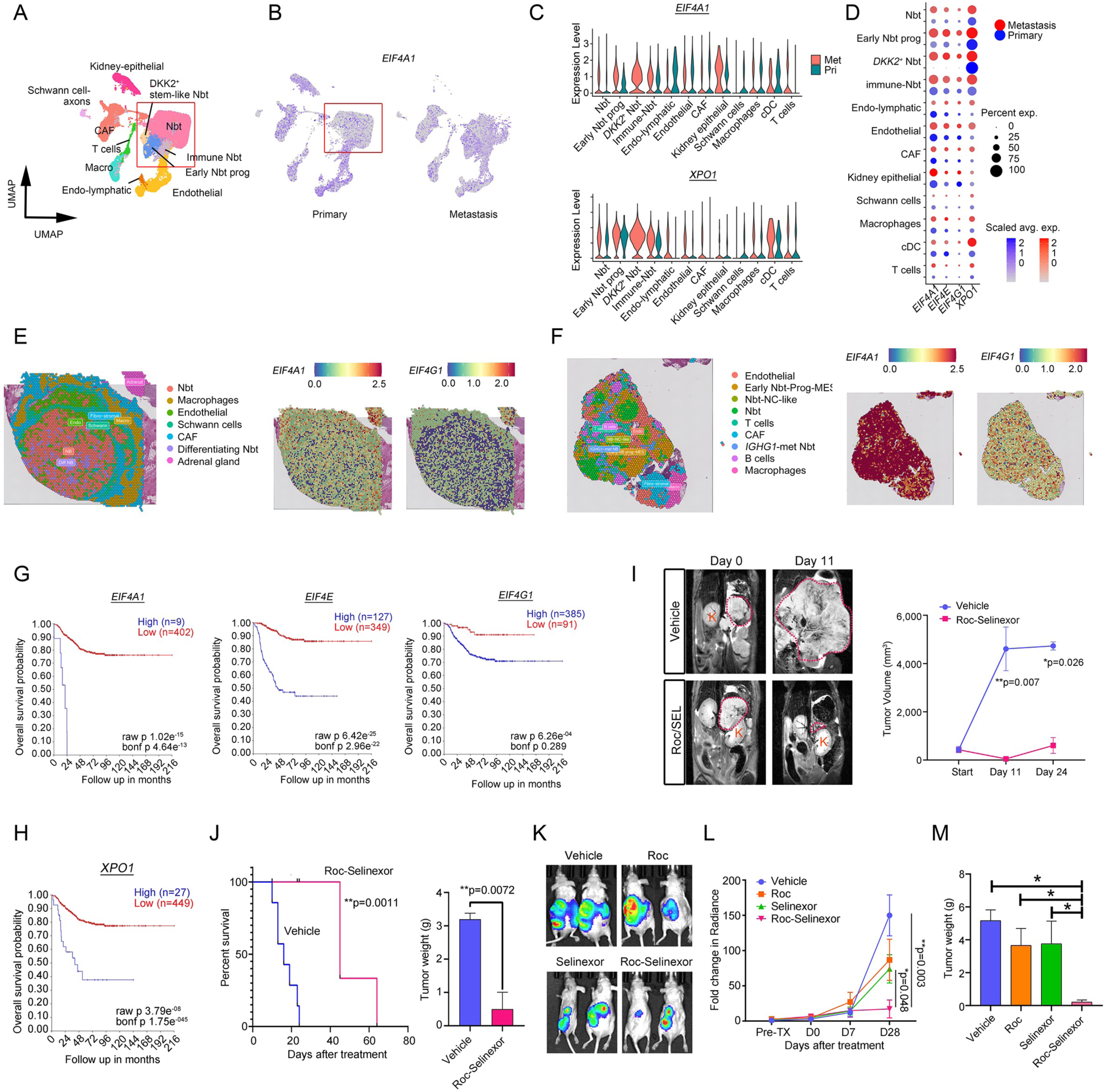
Dual targeting protein translation initiation and nuclear export halts NB growth. a. UMAP visualization of single-cell transcriptomes of primary and metastatic tumors resected post-treatment. Colors represent assigned cell types within NB tumor TMEs. b. UMAP visualization of expression of *EIF4A1* in NB primary and metastatic tumors. c. Violin plots of *EIF4A1* (upper) and *XPO1* (lower) expression in NB cell clusters. d. Dot plot of percent of cells in indicated NB cell clusters from primary and metastatic lesions that express *EIF4A1*, *EIF4E*, *EIF4G1*, and *XPO1*. e-f. Spatial transcriptomics analyses of expression of *EIF4A1* and *EIF4G1* in sections from e) primary and f) metastatic tumors. g. Kaplan–Meier overall survival analysis of patients from the Kocak dataset ^61^ (GSE45547) based on expression of EIF4F complex components. h. Kaplan-Meier overall survival analysis of patients from the Kocak dataset based on expression of *XPO1*. i. Left: Representative magnetic resonance images of SVJ129 mice allografted with primary mouse TH-MYCN^−/−^ NB cells in the adrenal fat pad treated with vehicle or the combination of rocaglamide and selinexor at day 0 and day 11. Red dotted lines outline tumor areas. The letter ‘K’ denotes the kidney. Right: Tumor volumes determined by MRI over time in vehicle (n=3 for Days 0 and 11; n=2 for Day 24) or rocaglamide and selinexor-treated mice (right; n=3 for all time points). \**P* < 0.05 and \*\**P* < 0.01, multiple t tests using the Holm-Sidak method. Data are means ± SEM. j. Left: Kaplan-Meier survival curves for TH-MYCN^−/−^ NB tumor-bearing SVJ129 mice treated with vehicle (n= 7) or the rocaglamide (Roc) and selinexor combination (n=6). *P* = 0.0011, log-rank tests. Right: Tumor weights of TH-MYCN^−/−^ allografts from mice treated with vehicle (n=3) or the combination of Roc and selinexor (n=3) at Day 24. \*\**P* < 0.01. Data are means ± SEM. k. Representative bioluminescence images of athymic nude mice xenografted with luciferase-expressing CHLA15 (CHLA15-Luc) engineered from a primary NB tumor from a high-risk patient and treated with vehicle, Roc, selinexor, or their combination. l. Quantification of bioluminescence signals as a function of time for CHLA15-Luc xenografted mice treated with Roc, selinexor, or their combination over time. Data are the means ± SEM (n > 5 mice). m. Tumor weights from CHLA15-Luc xenografted mice treated with Roc, selinexor, or the combination at Day 28. \**P* < 0.05, one-way ANOVA with Tukey’s multiple comparisons test. Data are means ± SEM.

To determine if targeting the translation initiation machinery in NBs halts tumor growth, we evaluated the antitumor efficacies of two eIF4A inhibitors, rocaglamide and didesmethylrocaglamide ^62^. *In vitro*, both inhibited growth of NB cell lines (Supplementary Fig. S9B-C). We next sought to identify signaling pathways that could be targeted to synergize with eIF4A inhibition. We discovered that expression of the gene encoding the nuclear export protein XPO1, which shuttles eIF4E to the cytoplasm, was substantially elevated in NB tumor cell populations (Fig. 8D) and that upregulated *XPO1* expression correlated with poor patient survival (Fig. 8H). *XPO1* overexpression is associated with cancer progression, treatment resistance, and inferior overall or progression-free survival ^63^. A recent study indicated that combined blockade of protein translation and nuclear export mediated by XPO1 enhances the anti-tumor activity relative to either inhibitor alone in myeloma cell lines ^64^. We found that XPO1 inhibition with the FDA-approved inhibitor selinexor synergized with the eIF4A inhibitor rocaglamide to suppress NB cell growth *in vitro* (Supplementary Fig. S9D-F). Notably, combining selinexor with rocaglamide synergistically inhibited NB tumor growth and prolonged survival in both the orthotopic allograft (Fig. 8I-J) and patient-derived xenograft models (Fig. 8K-M). Thus, our results indicate that dual targeting of the protein translation machinery and nuclear export may be a rational therapeutic strategy to treat high-risk NB.

## Discussion

NB is the leading cause of extracranial cancer-related deaths in children. Once NBs have metastasized, patients have poor prognosis despite aggressive multimodal therapy ^65,66^. Primary NB tumors have been extensively studied ^11–16^. Here, we characterized the unique cellular heterogeneity and transcriptional plasticity of lymph node NB metastases. By integrating single-cell multi-omics and spatial transcriptomics, we discovered that lymph node metastases harbored diverse tumor cell states, including mesenchymal-like and neural-crest stem-like cell phenotypes, that were less prevalent in primary tumors. This contrasts with the adrenergic differentiation signature that predominantly characterizes tumor cells in primary adrenal masses ^11–16^, reflecting their sympathoadrenal lineage origin. These findings support a model in which NB metastases adopt less differentiated, more plastic states conducive to invasion and therapy resistance ^9,10^. Importantly, lymph node metastases were enriched in EMT-related gene programs and transcriptional circuits such as those driven by ZEB1, TWIST1, and TGIF2, which are known to fuel the mesenchymal shift in tumor cell states facilitating dissemination and survival in secondary sites ^67,68^. We observed differences in chromatin accessibility and upregulated gene regulatory networks in metastatic cells compared to primary tumor cells. Genes upregulated in metastatic tumor cells are associated with early neural crest development and EMT. The mesenchymal prevalence in lymph node metastases differs from the adrenergic commitment of tumor cells in bone marrow metastasis and may reflect niche-specific pressures mediated by tumor milieu interactions at distinct metastatic sites^13,69^

Unlike primary NB masses, lymph node metastases have a pronounced immunosuppressive landscape enriched with immunosuppressive myeloid subsets, particularly pro-tumorigenic M2-like macrophages, and increased expression of immune checkpoint molecules such as CD47 and CD24. These features likely enable tumor cells to evade macrophage-mediated phagocytosis and foster a pro-tumorigenic niche ^70,71^. Immunosuppressive myeloid populations were also detected in NB bone marrow metastases ^69^, suggesting a conserved mechanism of immune evasion across metastatic sites. Additionally, metastatic sites showed reduced proportions of activated CD8^+^ T cells and NK cells, higher levels of expression of exhaustion markers (e.g., *TOX*, *NFAT5*), and upregulated TGFβ signaling, which has been reported to contribute to lymphocyte dysfunction and immune escape ^72,73^.

By integrating spatial transcriptomics with single-cell transcriptomics, we further delineated the cellular architecture of primary and metastatic NBs, revealing distinct tumor, immune, and stromal compartmentalization. In primary tumors, neuroblast clusters expressing adrenergic markers (e.g., *NPY* and *DBH*) dominated the tumor core and were bordered by stromal and immune components, with limited T cell infiltration. The lymph node metastases had more heterogeneous tumor architecture, with tumor cells located in closer proximity to lymphocytes than observed in primary tumors. In primary NBs, we detected neuroblast-derived paracrine signaling through MK and SPP1, factors that drive angiogenesis and tumor progression. In contrast, lymph node metastases displayed a unique tumor cell phenotype marked by IGHG1 expression, and the microenvironment was immunosuppressive resulting from activation of the TGFβ signaling pathway, potentially inducing immune exhaustion ^74^. The identification of active IGF signaling in metastatic NBs underscores key mechanisms facilitating EMT and tumor dissemination. Spatial transcriptomics further revealed tumor-immune cell interactions mediated by TGFβ signaling. The immunosuppressive and dysfunctional TME likely results in the lack of response of metastatic NBs to immune-based therapies. Thus, integrated single cell and spatial transcriptomics analysis uncover critical spatially defined TME interactions and targetable paracrine signals that contribute to NB metastasis, highlighting the importance of context-specific therapeutic strategies in high-risk NBs.

Integration of single-cell and spatial transcriptomics also revealed that there was robust activation of the eIF4F complex in metastatic NB tumor cell populations, highlighting the translational machinery as a crucial oncogenic driver. MYCN-driven high-risk NBs may enhance the eIF4F translation machinery to sustain tumor growth and metastasis, and elevated expression of eIF4A1, eIF4E1, and eIF4G1 correlate with poor prognosis. Pharmacological inhibition of eIF4A with rocaglamide or didesmethylrocaglamide effectively suppressed NB cell proliferation *in vitro* and significantly reduced tumor burden *in vivo*. Dysregulation of translation initiation leads to increased protein synthesis specifically of growth-related factors ^75,76^. Targeting eIF4E and other components of the translation machinery has shown potential in clinical trials in acute myeloid leukemia patients ^77,78^. Our findings underscore the translational machinery as a key vulnerability in aggressive NBs.

Furthermore, we identified upregulation of *XPO1* in metastatic tumors. XPO1 facilitates the nuclear export of critical tumor suppressors and oncoproteins, its upregulation is a hallmark of tumorigenesis, and its expression correlates with poor survival and aggressive disease ^79^. We showed that combining eIF4A inhibition with the FDA-approved XPO1 inhibitor selinexor resulted in a potent synergistic effect in cell culture and in profound tumor growth suppression and extended survival in both orthotopic and patient-derived xenograft models. Our findings demonstrate the therapeutic potential of combined inhibition of protein translation and nuclear export mechanisms in metastatic NBs.

In summary, our comprehensive characterization of cellular diversity and signaling networks in metastatic NBs provides a blueprint for further exploration of metastatic site-specific vulnerabilities. Our findings emphasize the need for therapies that target not only tumor-intrinsic mechanisms but also the immunosuppressive TME. Moreover, the integration of single-cell and spatial transcriptomics proved invaluable in elucidating the intricate interplay between tumor and microenvironmental components. By uncovering unique phenotypes and regulatory networks in lymph node metastases of NBs, we identified key mechanisms driving metastasis and therapy resistance. Our findings suggest that dual targeting of eIF4F-mediated translation initiation and XPO1-mediated nuclear export has potential to combat high-risk NB, particularly in patients with *MYCN*-amplified metastatic tumors, who currently have limited treatment options.

## Acknowledgements

The authors would like to thank Katie A. Belcher, Justin Ferrell, Xiang (Sean) Zhang, Andrew Potter and Hung-chi Liang for technical support. This study was funded in part by the grants from the Department of Defense W81WXH2010443 to QRL and LMNW, National Institute of Health (R01NS140460 and R01NS142389) to QRL and CancerFree KIDs to LSC.

## METHOD DETAILS

### Human Tumor Samples

All human patient samples were obtained with consent under approval and oversight by the Institutional Review Board (IRB) committees of Cincinnati Children’s Hospital Medical Center. Post-induction therapy NBs for single-cell studies were obtained from Cincinnati Children’s Hospital Medical Center Oncology Tissue Repository. Single-cell transcriptomics datasets of treatment-naïve high-risk NBs at adrenal gland (SCPCA sample ID: SCPCS000113 and SCPCS000106) or mediastinal mass (SCPCS000102) were obtained from Alex’s Lemonade Stand Single-cell Pediatric Cancer Atlas Portal https://www.biorxiv.org/content/10.1101/2024.01.07.574538v2 ^80,81^.

### Animals

All studies complied with all relevant animal use guidelines and ethical regulations. All animal use and study protocols were approved by the Institutional Animal Care and Use Committee (IACUC) at the Cincinnati Children’s Hospital Medical Center, Ohio, USA. TH-MYCN transgenic mice (TH-MYCN+/−) on SVJ129 background overexpress MYCN proto-oncogene in the sympathoadrenal lineage and develop spontaneous, aggressive adrenal NB and were a kind gift from Dr. William A. Weiss ^91^.

### Mouse Orthotopic Allografted NB Model

Primary mouse NB isolated from TH-MYCN+/− mice were minced on a plate and enzymatically digested with collagenase IV (2 mg/ml; Gibco) at 37°C with agitation on a gentleMACS Octo Dissociator using the NTDK program. After digestion, samples were filtered through a 40-µm cell strainer, washed with RPMI containing 10% fetal bovine serum (FBS), and centrifuged for 5 min at 500*g*. Red blood cells and debris were removed by red blood cell lysis buffer (Roche) according to the manufacturer’s specifications. Pelleted cells were then resuspended in RPMI with FBS, filtered through a 35-µm cell strainer, and assessed for viability using trypan blue. 3 × 10^5^ cells resuspended in 15 µl Matrigel (Corning) and DMEM/10% FBS at 1:1 ratio were injected into the left adrenal gland of 6-8 weeks old SVJ129 mice using a Hamilton syringe fitted with a 27G needle.

Orthotopically engrafted mouse TH-MYCN+/− NB were imaged using a 7-Tesla small animal magnetic resonance imaging (MRI) at the CCHMC Small Animal Imaging Core. Tumor volumes were measured by T2-weighted coronal and transverse images acquired from contiguous 1 mm thick slices through the mouse abdomen. Tumor-bearing TH-MYCN+/− mice (tumors > 400 mm^3^) were randomly assigned to two treatment groups: vehicle and rocaglamide (Roc)/Selinexor combination (n ≥ 4 per group). For combination treatment of Roc and Selinexor, mice were administered 3mg/kg Roc dissolved in 30% 2-Hydroxypropyl-beta-cyclodextrin (HPBCD), via IP injections every other day and 6.66 mg/kg Selinexor dissolved in 0.6% Polyvinylpyrrolidone (PVP K-30) and 0.6% Pluronic F-68, via oral gavage (PO) every other day. For Roc administration, mice received a lead-in dose of 1mg/kg Roc for the first dose, 1.5mg/kg for the second dose, and 2 mg/kg for the third dose, all via IP injections. Mice in the vehicle group were administered HPBCD: PVP K-30 and Pluronic F-68 at 1:1 ratio via oral gavage. Mice were monitored for body weight loss and survival, and tumor volume assessed via T2 weighted MRI imaging bi-weekly. Tumor weight was measured at endpoint upon tumor dissection at tissue harvest.

### Patient Cell Line-derived Xenograft Models and Bioluminescence Imaging

Patient-derived NB cell lines (MYCN non-amplified: CHLA15, CHLA20, MYCN-amplified: SK-N-BE2, COG-n-496 and COG-n-623) were obtained from Children’s Oncology Group (COG). CHLA15, CHLA20, COG-n-496 and COG-n-623 cells were cultured in Iscove’s Modified Dulbecco’s Medium (IMDM) supplemented with 20% Fetal Bovine Serum (FBS), 4 mM L-Glutamine, 1X ITS (5 µg/mL insulin, 5 µg/mL transferrin, 5 ng/mL selenous acid; Fisher Scientific). SK-N-BE2 cells were grown in RPMI-1640 supplemented with 10% Fetal Bovine Serum and 2mM L-Glutamine. Luciferase-expressing NB cells were generated by transducing NB cells with CMV-Firefly luciferase lentivirus (Cellomics Technology LLC). Puromycin-resistant clones were assessed for luciferase activity using the Luciferase Reporter Assay System (Promega), and the highest expressing clone of CHLA15 cells (CHLA15-Luc), CHLA20 cells (CHLA20-Luc), COG-n-623 (COG-n-623-Luc) were selected for *in vivo* grafting.

For orthotopic cell engraftment in the adrenal gland, 5 × 10^5^ cells CHLA15-Luc, CHLA20-Luc or COG-n-623-Luc cells resuspended in 15 µl Matrigel (Corning) and IMDM/20% FBS at 1:1 ratio were injected into the left adrenal gland of 6-10 weeks old athymic nude mice, using a Hamilton syringe fitted with a 27G needle. Following tumor establishment, mice were randomized into 4 groups: vehicle, Roc alone, Selinexor alone, and Roc/Selinexor combination (*n* ≥ 5 each). For single Roc treatment, mice were administered 3mg/kg Roc dissolved in 30% 2-Hydroxypropyl-beta-cyclodextrin (HPBCD), via IP injections every other day, with a lead-in dose of 1mg/kg Roc for the first dose, 1.5mg/kg for the second dose, and 2 mg/kg for the third dose. For single Selinexor treatment, mice were administered 10 mg/kg Selinexor via oral gavage (PO) every other day. Tumor growth was monitored weekly by bioluminescence imaging (BLI) using a Xenogen IVIS Spectrum (Caliper). Data acquisition and analysis was performed using the Living Image® software (Caliper LS).

### Preparation of Single-Nuclei Suspensions for 10X scMultiome assay

Snap-frozen human NBL specimens were lysed in ice-cold nuclei lysis buffer (Nuclei Extraction Buffer (Miltenyi Biotec) containing DTT (Sigma-Aldrich) and RNAse inhibitor (Sigma-Aldrich)) for 5 min on a gentleMACS Octo Dissociator at 4°C with agitation. After lysis, samples were filtered through a 70-µm cell strainer, washed with lysis buffer and centrifuged for 5 min at 500 × g at 4°C. Nuclei pellet was resuspended with ice-cold 0.1% BSA in PBS with RNAse inhibitor, filtered through a 30-µm cell strainer and centrifuged at 500 × g at 4°C for 5 min. Next, nuclei were permeabilized by resuspending the pellet in 0.1x lysis buffer (10 mM NaCl, 10 mM Tris 7.4, 3 mM MgCl2, 0.1% IGEPAL CA-63, 0.01% Digitonin, 1% BSA, 1 mM DTT, 1 U/µL of Protector RNase inhibitor (Sigma)) and incubated on ice for 2 min. 1 mL Wash Buffer (10 mM NaCl, 10 mM Tris 7.4, 3 mM MgCl2, 0.1% Tween-20, 1% BSA, 1 mM DTT, 1 U/µL of Protector RNase inhibitor (Sigma)) was added. The nuclei were centrifuged at 500 x g for 5 min and the supernatant was discarded. The nuclei pellet was resuspended in diluted nuclei buffer (1x Nuclei buffer Multiome kit (10x Genomics)), 1 mM DTT, 1 U/µL of Protector RNase inhibitor (Sigma) according to the 10X Genomics protocol for Nuclei Isolation from Complex Tissues for Single Cell Multiome ATAC + Gene Expression Sequencing. Nuclei integrity was assessed using Trypan blue and nuclei concentration was adjusted for 10X Genomics at CCHMC Single Cell Genomics Facility (Cincinnati, OH) using a manual hemocytometer to achieve a targeted nuclei recovery of 3,000-10,000 per sample.

### Chromium Single Cell Multiome ATAC + Gene Expression Sequencing

Single-nuclei libraries were generated using the 10x Chromium Single-Cell Instrument and NextGEM Single Cell Multiome ATAC + Gene Expression kit (10x Genomics) according to the manufacturer’s protocol. Briefly, single patient NB nuclei were incubated for 60 min at 37°C with a transposase that fragments the DNA in open regions of the chromatin and adds adapter sequences to the ends of the DNA fragments. Next, transposed nuclei were combined with Single Cell Multiome ATAC + Gene Expression Gel Beads to generate nanoliter-scale gel bead-in-emulsions (GEMs) on a Chromium Next GEM Chip J. GEMs were incubated in a C1000 Touch Thermal Cycler (Bio-Rad) under the following program: 37°C for 45 min, 25°C for 30 min and hold at 4°C to produce 10x barcoded DNA from the transposed DNA (for ATAC) and 10x barcoded, full-length cDNA from poly-adenylated mRNA (for GEX). Next quenching reagent (Multiome 10x kit) was used to stop the reaction. After quenching, single-cell droplets were dissolved, and the transposed DNA and full-length cDNA were purified using Cleanup Mix containing Silane Dynabeads.

To fill gaps and generate sufficient mass for library construction, the transposed DNA and cDNA were amplified via PCR: 72°C for 5 min; 98°C for 3 min; 7 cycles of 98°C for 20 s, 63°C for 30 s, 72°C for 1 min; 72°C for 1 min; and hold at 4°C. The pre-amplified product was used as input for both ATAC library construction and cDNA amplification for gene expression library construction. Illumina P7 sequence and a sample index were added to the single-strand DNA during ATAC library construction via PCR: 98°C for 45 s; 7-8 cycles of 98°C for 20 s, 67°C for 30 s, 72°C for 20 s; 72°C for 1 min; and hold at 4°C. The sequencing-ready ATAC-seq library was cleaned up with SPRIselect beads (Beckman Coulter). Barcoded, full-length pre-amplified cDNA was further amplified via PCR: 98°C for 3 min; 6-9 cycles of 98°C for 15 s, 63°C for 20 s, 72°C for 1 min; 72°C for 1 min; and hold at 4°C. Subsequently, the amplified cDNA was fragmented, end-repaired, A-tailed and index adaptor ligated, with SPRIselect cleanup in between steps. The final gene expression library was amplified by PCR: 98°C for 45 s; 5-16 cycles of 98°C for 20 s, 54°C for 30 s, 72°C for 20 s. 72°C for 1 min; and hold at 4°C. The sequencing-ready GEX library was cleaned up with SPRIselect beads. Before sequencing, the fragment size of every library was analyzed using the Bioanalyzer high-sensitivity chip. All 10x Multiome ATAC libraries were sequenced on NovaSeq6000 (Illumina) with the following sequencing parameters: 50 bp read 1 – 8 bp index 1 (i7) – 16 bp index 2 (i5) - 49 bp read 2. All 10x Multiome GEX libraries were sequenced on NovaSeq6000 instruments with the following sequencing parameters: 28 bp read 1 – 10 bp index 1 (i7) – 10 bp index 2 (i5) – 75 bp read 2. Libraries were sequenced to a target depth of 250 million read pairs per sample.

### scRNA- and scATAC-seq Data Preprocessing and Quality Control

Demultiplexed scRNA- and scATAC-seq fastq files were input into the Cell Ranger ARC pipeline (version 2.0.0) from 10x Genomics to align reads to the hg38 human reference sequence and generate barcoded count matrices of gene expression and ATAC data. scRNA-seq and snMultiome-seq data was analyzed with the open-source Seurat ^82^ and ArchR packages ^41^ implemented in the R computing environment.

### Dimensionality Reduction and Clustering

For each snRNA dataset, unsupervised clustering was performed using R package Seurat (v4.4.0) ^82^. Based on the distribution of cells ordered by percentage of mitochondrial genes and detected gene numbers, we excluded those cells with either more than 9,000 detected genes or less than 200, and mitochondrial content of more than 20%. FindVariableFeatures was set at nfeatures = 5,000. We used the filtered expression matrix to identify cell subsets. The filtered gene expression matrix was then normalized using Seurat’s NormalizeData function, in which the number of UMIs of each gene was divided by the sum of the total UMIs per cell, multiplied by a scale factor of 10,000, and then transformed to logscale (ln (UMI-per-10000+1)). Highly variable genes were identified based on overdispersion of genes in each gene group binned with aggregate expressions, and used for Principal component analysis (PCA). We then performed clustering using graph-based clustering and visualized using Uniform Manifold Approximation and Projection (UMAP) with Seurat function RunUMAP.

To identity cell type in tumor sample datasets, we input marker gene lists generated by Seurat FindMarkers function in ToppGene ^83^ to identify the top cell-type makers or cell identities. Differentially expressed genes (DEGs) in a given cell type compared with all other cell types were determined with the FindAllMarkers function from the Seurat package (one-tailed Wilcoxon rank sum test, p values adjusted for multiple testing using the Bonferroni correction). For computing DEGs, all genes were probed if they were expressed in at least 25% of cells in either of the two populations compared and the expression difference on a natural log scale was at least 0.25. The “scaled average expression” values for each gene in dot plots are scaled by subtracting mean expression of the gene and dividing by its SD.

Gene module scores for activation and exhaustion gene expression programs defined by T cell activation and exhaustion gene regulators in lymphocytes were generated using AddModuleScore function in Seurat. M1 and M2 marker genes from Xue et al., ^42^ were used as reference gene sets for computing gene module scores across myeloid cells in NB samples.

### Batch Correcting and Multiple Datasets Integration

We used the scRNA-seq integration platform on Seurat ^84^ to correct for technical differences between datasets (i.e. batch effect correction) and to perform comparative scRNA-seq analysis within the same tumor type from multiple datasets and between different tumor types. These methods identify cross-dataset pairs of cells that are in a matched biological state termed ‘anchors’ for batch effect removal and multiple dataset integration.

### Deconvolution Analysis

CIBERSORTx ^35^ was used to perform deconvolution analysis of NB scRNA-seq against signature gene input of developing lineages in sympathoadrenal regions during human embryogenesis from post-conception weeks 6-14 (GSE147821) ^15^. The signature gene input was generated by differential gene analysis using Altanalyze ^85^ of the GSE147821 dataset.

### Inferred CNV Analysis from scRNA-seq

Malignant cells were identified by inferring large-scale chromosomal copy-number variations (CNVs) in each single cell based on a moving averaged expression profiles across chromosomal intervals by inferCNV ^30,86^. We combined CNV classification with transcriptomic-based clustering and expression of non-malignant marker genes to identify malignant and non-malignant cells. Non-malignant cells displayed high expression of specific marker genes and no apparent CNVs. These include macrophages, T cells, fibroblasts, endothelial cells etc. We analyzed post-induction therapy post-induction therapy high-risk NBL at primary sites from integration of 6 patient scRNA-seq datasets and NB metastases from integration of 5 datasets.

### Inferring Developmental Potential Using scRNA-seq Data by FitDevo

Developmental potential of NB-derived lineage populations within high-risk NBLs at adrenal gland and lymph node metastases was inferred by FitDevo via calculating the correlation between sample-specific gene weight SSGW and gene expression ^24^.

### Pseudotime Trajectory Analysis and Differentiation Potential Evaluation

Slingshot ^25^ is a top performer for cellular trajectory from single-cell RNA-seq data, according to a benchmarking study reported by ^87^. We used Slingshot to model the pseudotime trajectory of NB-derived tumor populations between post-induction therapy high-risk NBL at primary and metastatic locations. The input matrix was filtered and normalized by the R package Seurat and cell types were annotated and provided as labels for Slingshot. We did not provide any further prior information about origin and end cell types of trajectories. In addition, we used the R package CytoTRACE v.0.3.3 to predict differentiation status of NB lineage cells from clinical scRNA-seq data ^88^.

### Novel Expression Programs of Intra-tumoral Heterogeneity by NMF

We applied non-negative matrix factorization (NMF) to extract transcriptional programs of NB-derived tumor populations from relative expression data (with conversion of negative values to zero) as previously described^36^. We used scater implementation of runNMF for the analysis. We set the number of components to 30 for each dataset (ncomponents=30, ntop=5000). For each of the resulting factors, we considered the top 100 genes with the highest NMF scores from each resulting NMF factor as characteristics of that given factor. All single cells of the NB lineage populations were then scored according to these NMF programs. NMF programs were clustered with hierarchical clustering on the scores for each program. This revealed five correlated sets of programs for the distinct NB-derived clusters in primary NB and six in NB metastasis analysis. We used pheatmap function in the R package to generate the heatmap of signature expression (log (NMF loading+1)) with cells sorted by NMF gene modules grouped by key meta-programs with annotation of select gene markers in NB at primary and metastatic locations.

### Gene Regulatory and TF Network and scATAC-Seq Data Analysis

To characterize underlying gene regulatory network and infer transcription factor activities for NB-derived tumor populations in our scRNA-seq dataset, we used the single-cell regulatory network inference and clustering (SCENIC) package to identify gene regulatory modules and retain those with a cis-regulatory binding motif for upstream regulators ^37^. By GENIE3, we estimated co-expression modules between TFs and putative target genes, followed by cis-regulatory motif analysis using RcisTarget and pruning of indirect targets lacking motif binding site. By AUCell, the resulting regulatory module (regulons, modules of target genes co-expressed with TFs and enriched with motifs for correct upstream regulators) activities in each cell were then binarized. The input matrix was the normalized expression matrix of cells of interest. TF regulatory networks (regulon, *x* axis) derived by SCENIC for each NB-derived tumor subpopulation was plotted against their normalized specificity score (*y* axis).

ArchR ^41^ was used for all scATAC-seq analyses according to default parameters including quality control and cell filtering, dimension reduction, genome browser visualization, gene expression data preprocessing and cell annotation, DNA accessibility data processing, joint data visualization, differential accessibility and motif enrichment. For each sample, count matrices were loaded in ArchR and selected for barcodes that appeared in both the scRNA-seq and scATAC-seq datasets. Samples in ArchR were quality control filtered for nuclei with 200-50,000 RNA transcripts, <1% mitochondrial reads, <5% ribosomal reads, TSS enrichment >6, and >2,500 ATAC fragments. Quality control filtered nuclei subsequently underwent automated removal of doublets using the filterDoublets function in ArchR, which identifies and removes the nearest neighbors of simulated doublets.

### Spatial Transcriptomics

Spatial transcriptomics of NB at primary adrenal gland and metastatic lymph nodes was performed using the Visium CytAssist-enabled workflow (10x Genomics) according to manufacturer’s instructions. Formalin-fixed paraffin-embedded (FFPE) NB specimens were sectioned at 10 µm using a microtome on a clean glass slide. NB section was H&E stained and imaged by the CCHMC Single Cell Genomics Facility using the Keyence BZ-X810. Once histological information was recorded, the gene expression information was acquired. Visium Human Transcriptome Probe Set v2.0 (consisting of probe pairs for each targeted gene) was subjected to the tissue enabling probe-pair hybridization and ligation. Post ligation, the plain glass slides and Visium slide were mounted into the CytAssist instrument, and the ligated probe-pairs were transferred from the tissue on the plain glass slide to the barcoded-array capture areas on the Visium slide maintaining spatial configurations (11 x 11mm or 6.5 x 6.5mm) to capture gene expression data. Subsequently, the Visium slide containing the probe-pairs was removed from the instrument, and the probes were extended to incorporate the spatial barcoding information. The barcoded products were released from the Visium slide, collected and further processed into sequencing libraries. Libraries were sequenced on NovaSeq6000 to a target depth of at least 25,000 read pairs per tissue covered spot on Capture Area of the Visium slide.

Visium FASTQ files were input into the Space Ranger pipeline (version 2.0.0) from 10x Genomics to generate barcoded count matrices of gene expression data with spatial information. Spatially-resolved RNA-seq data was analyzed using the Seurat toolkit Spatial Analysis Vignette ^89^. snRNA-seq data from the corresponding NB specimens were used as reference datasets for cell type annotation.

### Using Receptor-Ligand Pairs to Infer Cell-Cell Interactions

CellChat v2 ^53^ was used to infer the intercellular communication from spatially resolved transcriptomics datasets, for further data exploration, analysis, and visualization. We applied CellChat v2 to NB spatial transcriptomics data to systematically computes and categorizes communication probabilities of ligand-receptor pairs into functionally relevant signaling pathways. Using pattern recognition analysis, we identified conserved and distinct global communication patterns and major signaling pathways enriched for each cell type in NB at primary adrenal gland and lymph node metastasis.

### Total RNA Isolation, RNA-sequencing and Data Analysis

RNA of frozen post-induction therapy NB patient samples was extracted using TRIZOL (Life Technologies) followed by purification using a Direct-zol RNA MiniPrep Kit (Zymo Research). RNA-seq libraries were prepared and sequenced by an Illumina Novaseq6000 sequencer. All RNA-Seq data were aligned to hg19 using TopHat with default settings. We used Cuff-diff to (1) estimate FPKM values for known transcripts and (2) analyze differentially expressed transcripts. In all differential expression tests, a difference was considered significant if the q value was less than 0.05 (Cuff-diff default). Heatmap of gene expression was generated based on log2 [FPKM] by AltAnalyze (AltAnalyze.org) with normalization of rows relative to row mean ^85^. GO-analysis of gene expression changes was performed using Gene Set Enrichment (GSEA, http://www.broadinstitute.org/gsea/index.jsp). Normalized enrichment score (NES) reflects the degree to which the gene-set is overrepresented at the top or bottom of a ranked list of genes. The GSEA scatterplots showing upregulated and downregulated pathways were plotted according to (https://www.biostars.org/p/168044/). Genes categorized with negative or positive NES are downregulated or upregulated, respectively. Circle size is defined as the number of genes represented in the leading-edge subset, i.e., the subset of members within a gene set that shows statistically significant, concordant differences between two biological states and contribute most to the NES. Gene sets with FDR q values < 0.25 are plotted as a function of NES. Circle colors represent FDR q values. We used ToppCluster (https://toppcluster.cchmc.org/) to construct the network of genes belonging to over-represented GO-term categories. For the volcano plot, the up-regulated and down-regulated genes were represented by red or blue dots respectively (fold change >2, adj. p < 0.1 between NBL at primary and metastatic locations). Gray dots represent insignificantly changed genes with p > 0.1. The gene expression signatures for pathway analysis were from Molecular Signatures Database v5.1. The mean expression values were calculated using all genes within a given signature for the heatmap analysis ^90^.

### Immunohistochemistry (IHC)

For paraffin processing and embedding, normal human peripheral nerves and NB patient samples at primary adrenal gland and metastatic lymph nodes were dissected, perfused and fixed overnight in 4% PFA, embedded in paraffin and sectioned at 5 µm. For paraffin sections, slides were deparaffinized, rehydrated and subjected to citrate-based or Tris-EDTA antigen retrieval. Endogenous peroxidase activity was blocked by incubation with hydrogen peroxide. Sections were incubated overnight with primary antibodies. Following three washes with PBS, sections were incubated for 1 h with biotinylated secondary antibodies, followed by ABC kit (Vector labs) application and the peroxidase/diaminobenzidine (DAB) method to visualize signals under light microscopy.

For immunoperoxidase staining, we used antibodies to IBA1 (Rabbit; Wako Chemicals, 019-19741), CD3 (Rat; Biorad, MCA1477), CD163 (Rabbit; Proteintech, 166461AP), eIF4AI-II (#sc-50354) and eIF4G (#sc-133155) (Santa Cruz) and eIF4E (Abcam #ab33768). All immunoperoxidase-labeled images were acquired on an Olympus BX53 brightfield microscope. For quantification of IBA1^+^ immunolabeled cells on coverslips, multiple images were taken from each coverslip to obtain representative images from all areas of the coverslip and at least 1000 cells/coverslip were counted using NIH ImageJ software.

### *In vitro* Compound Treatment and Cell Proliferation Assays

For NB cell lines CHLA15, CHLA20, SK-N-BE2 and COG-n-496, dose–response testing was performed in a 96-well format using Premix WST-1 Cell Proliferation Assay System (Takara Bio) or resazurin proliferation assay. Briefly, ∼18-20 hours post-seeding, cells were treated with serial dilutions of Roc, DDR, and/or Selinexor in full growth medium and incubated for 72 hours. Cell viability was determined by reading WST-1 colorimetric and resazurin fluorescence assays on a SpectraMax® ABS Plus Microplate Reader (Molecular Devices). Dose–response curves were plotted using GraphPad Prism 8.0 software, and the drug concentrations inhibiting cell growth by 50% (IC_50_) relative to DMSO were determined using nonlinear regression (curve fit) analysis. Drug combination 2D and 3D dose-response matrices of Roc or DDR and Selinexor were plotted by SynergyFinder Plus ^92^.

### Quantification and Statistical Analysis

Statistical analysis was performed using R (version >=4.4) and GraphPad Prism 8. Significance values are described in figures, figure legends, and/or the results section. Alternatively, significance levels are indicated as asterisks, signifying as * p<0.05, ** p<0.01, *** p<0.001. *In vitro* cell viability data were calculated from at least three independent experiments, performed as duplicates for each condition. Data points represent mean values ± SEM, as specified in figure legends.

### Data and Software Availability

The high-throughput sequencing data that support the findings of this study have been deposited in the NCBI Gene Expression Omnibus (GEO) under accession numbers GSE309267, GSE309268, and GSE309269.

## Supplemental figure legends

**Supplementary Fig. S1.**
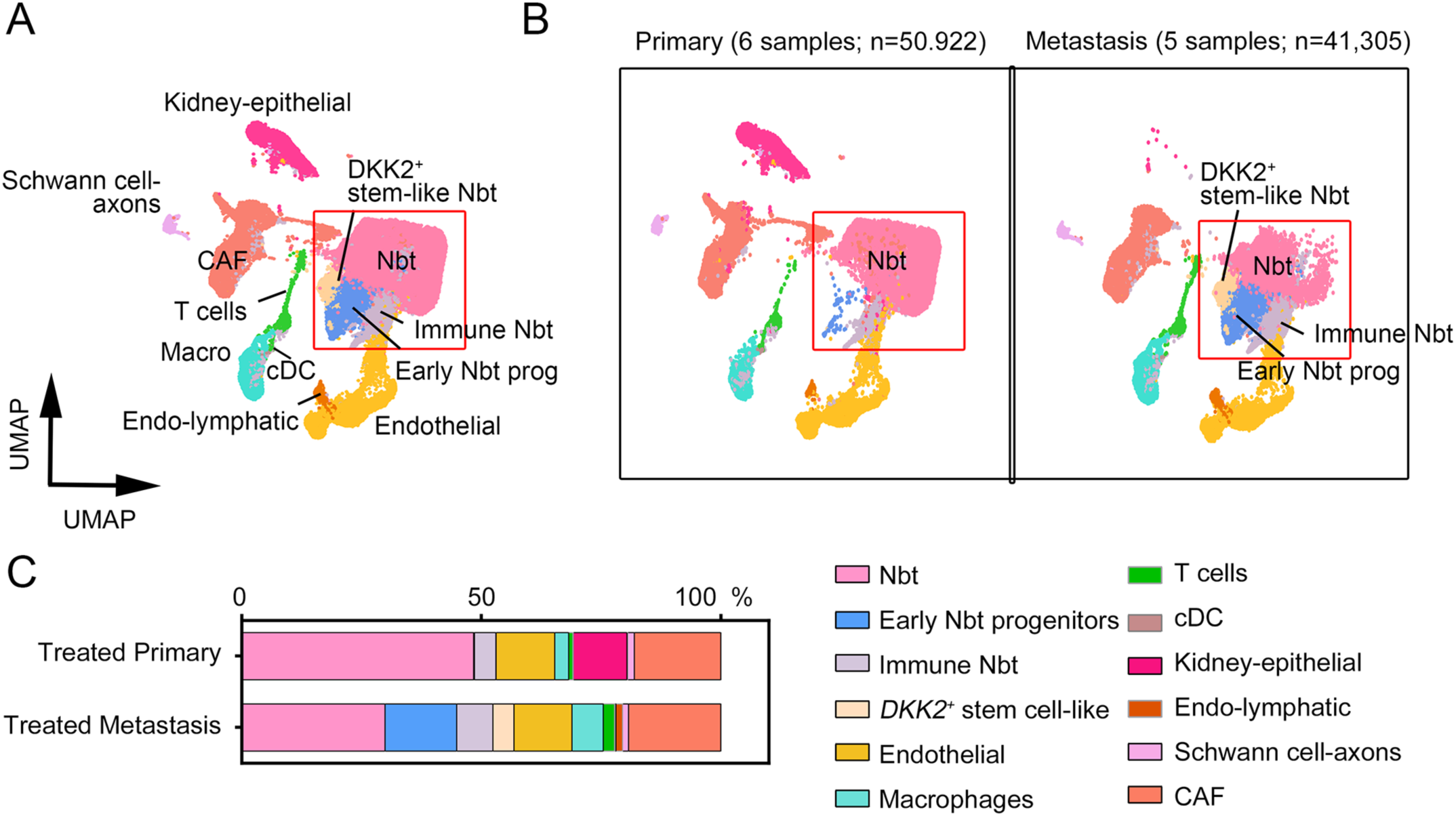
Comparison of cellular landscapes of high-risk primary NB and metastatic lesions resected post-treatment. a. UMAP visualization of single-cell transcriptome data of 92,227 cells from primary adrenal glands and lymph node metastatic sites of NBs from patients post-treatment. Colors represent assigned cell types within NB tumors. b. Side-by-side UMAP visualization of post-treatment NBs at adrenal glands (n=6; 50,922 cells) and at lymph node metastases (n=5; 41,305 cells). Red boxes denote tumor cell phenotypes in the NB tumor. c. A summary of cell type proportions in post-treatment NBs at primary and metastatic sites.

**Supplementary Fig. S2.**
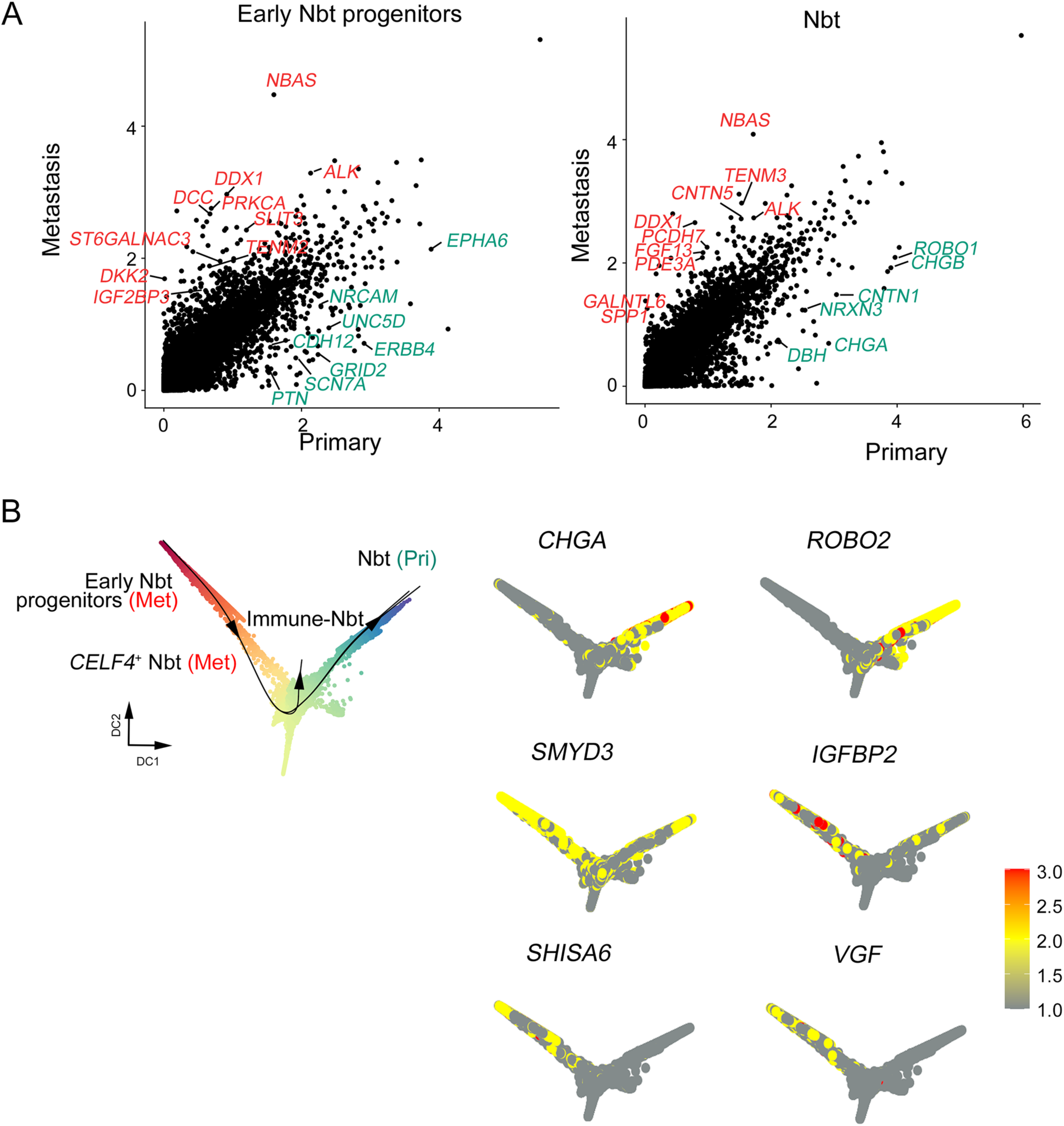
Differential gene marker expression in primary and lymph node metastatic sites. a. Differentially expressed genes of early NB progenitor populations (left) and tumor cells (right) from post-treatment NBs at primary and metastatic sites. Color labels indicate visual outliers on a scatter plot in primary and metastatic NBs. b. UMAP plots of marker genes within NB clusters based on Slingshot trajectories.

**Supplementary Fig. S3.**
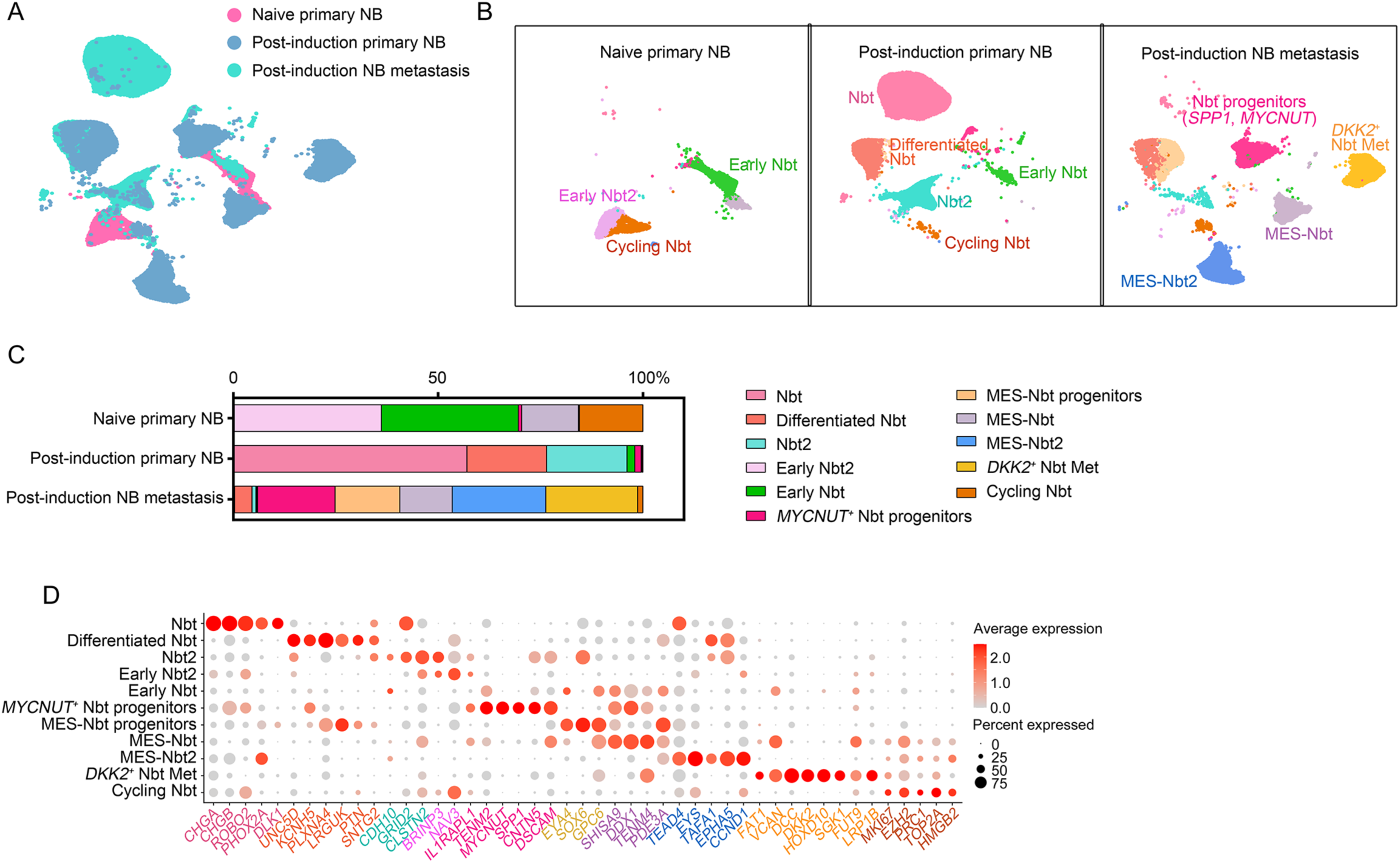
Cell states in NB tumors are heterogeneous. a. UMAP visualization of integrated scRNA-seq datasets of tumor cell phenotypes in primary tumors of treatment-naïve high-risk NB patients and in primary and metastatic tumors of post-treatment high-risk NB patients. Colors represent NB cell tumor types. b. UMAP visualizations of tumor cell subclusters (red rectangles in (Figure 1c) in primary tumors of treatment-naïve patients (8,905 cells), primary tumors of post-treatment patients (25,434 cells), and lymph node metastases of post-treatment patients (23,301 cells). Cells are colored according to tumor cell type. c. Proportions of NB-derived cell subtypes in primary tumors from treatment-naïve patients, primary tumors of post-treatment patients, and lymph node metastases of post-treatment patients. d. Marker gene expression in NB-derived cell clusters from NB tumors. The color represents scaled average expression of marker genes in each cell type, and the size indicates the proportion of cells that express marker genes.

**Supplementary Fig. S4.**
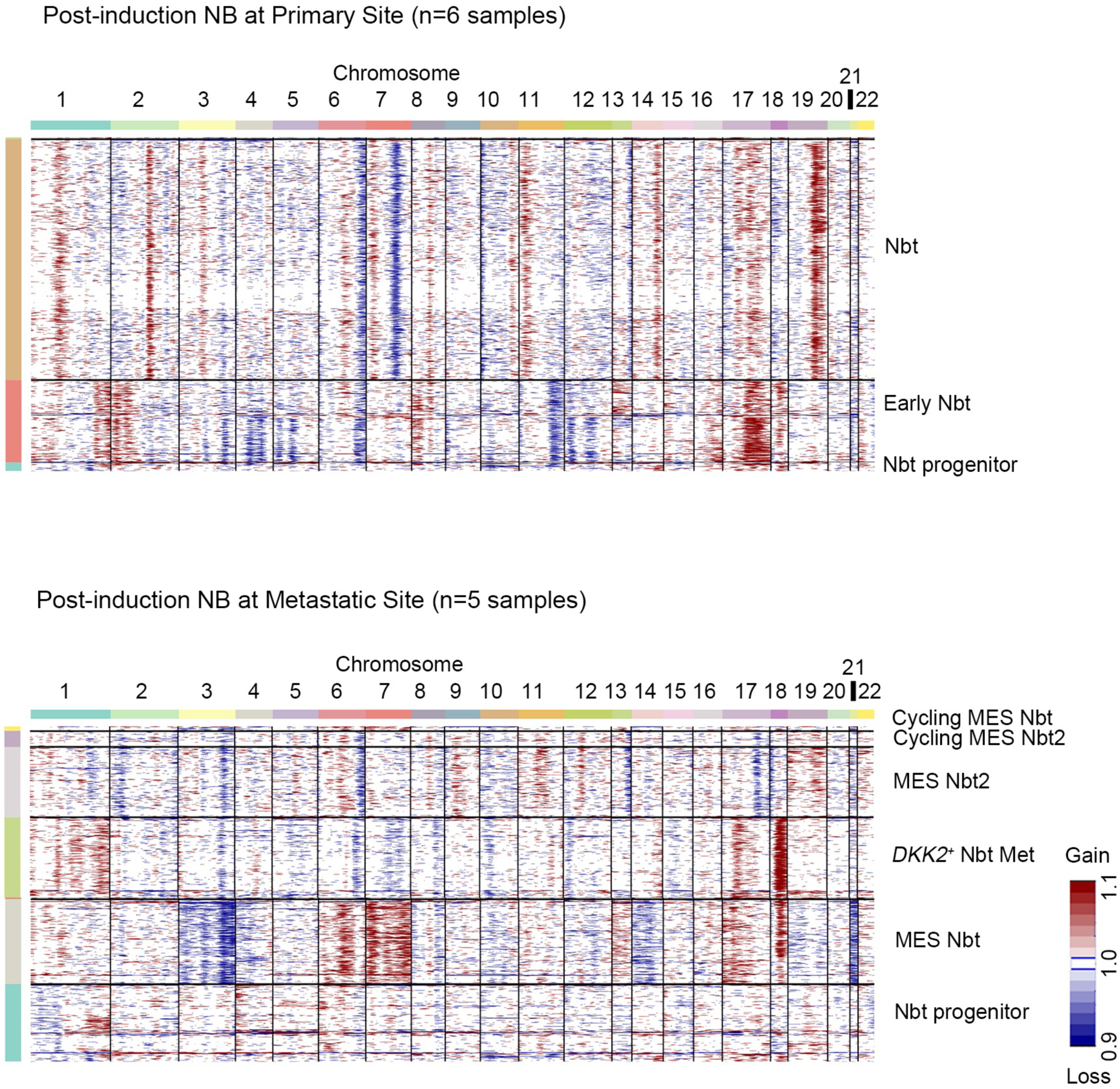
CNVs inferred from snRNA-seq of NB tumor cells from primary and metastatic lesions. CNVs detected in snRNA-seq data in post-treatment primary (n=6; left panel) and metastatic samples (n=5; right panel) with cut-off for the minimum average read counts per gene among reference stromal and immune cells set at 0.1. Each row corresponds to a cell, ordered by tumor cell phenotype, and clustered within each tumor by CNV patterns.

**Supplementary Fig. S5.**
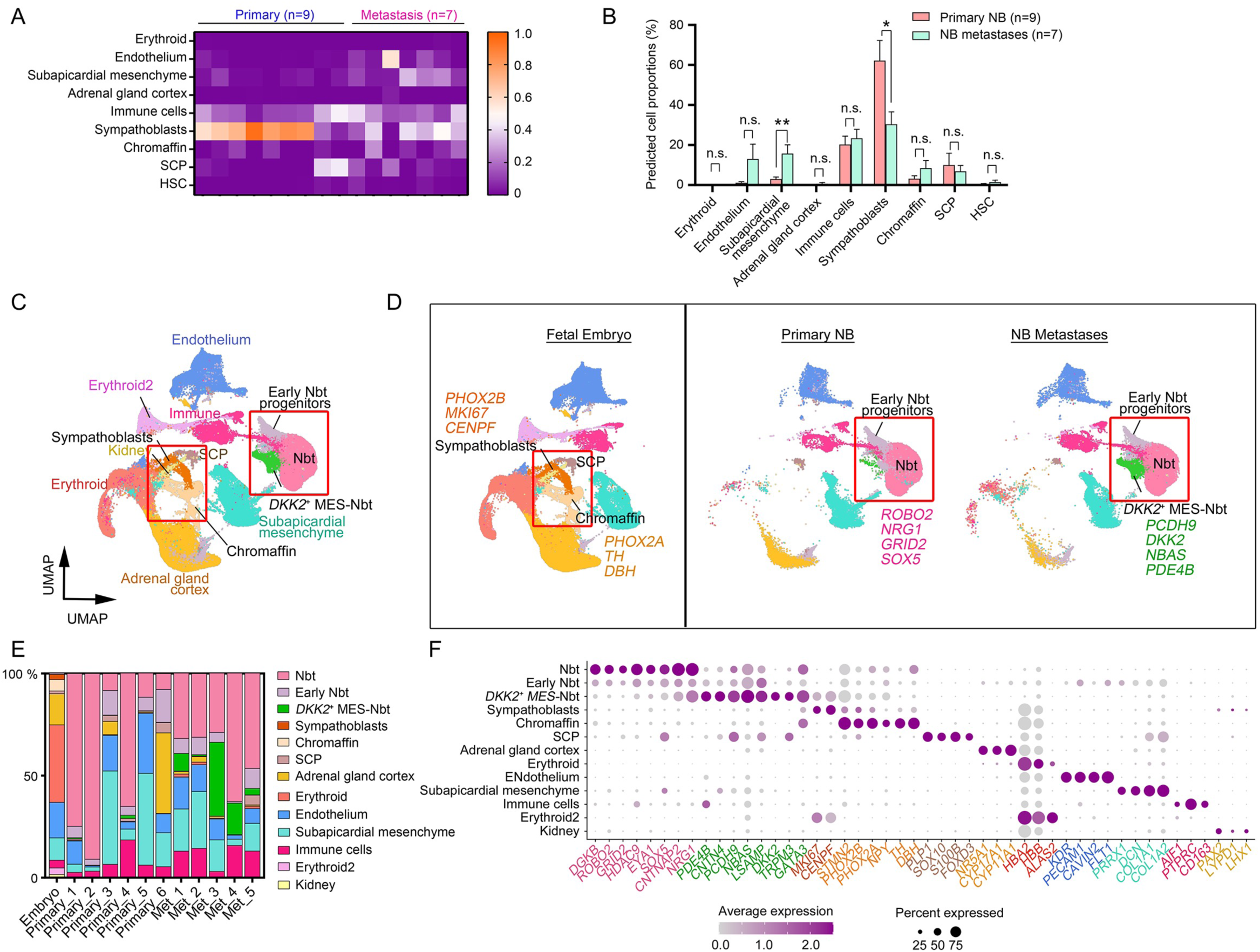
NB metastatic cell states have immune and mesenchymal signatures distinct from embryonic sympathoblasts. a. Heatmap of cell proportions in tumor cell states in post-treatment NBs at primary (n=9) and metastatic sites (n=7) relative to those in developing lineages in sympathoadrenal regions during human embryogenesis from post-conception weeks 6-14 (GSE147821) ^15^ determined using CIBERSORTx deconvolution of scRNA-seq data. b. Predicted proportions of indicated cell types in primary and metastatic NBs. *P < 0.05 and **P < 0.01, multiple t tests using the Holm-Sidak method. Data are means ± SEM. c. UMAP visualization of integrated scRNA-seq datasets of human embryogenesis lineages (GSE147821; n=134,491), post-treatment primary NB (n= 50,922 cells), and metastatic NB (n= 41,305 cells). d. Side-by-side UMAP visualizations of human embryogenesis lineages, post-treatment primary NB, and metastatic NB. Red rectangles denote the sympathoadrenal lineages in fetal embryo and NB-derived tumor cell phenotypes in NBs. e. Cell type proportions in human embryogenesis lineages (Embryo) and individual post-treatment primary and metastatic samples. f. Marker gene expression in embryogenesis lineages and NBs from primary and metastatic tumors. The color represents scaled average expression of marker genes in each cell type, and the size indicates the proportion of cells that express the marker gene. SCP, Schwann cell precursors; HSC, hematopoietic stem cells.

**Supplementary Fig. S6.**
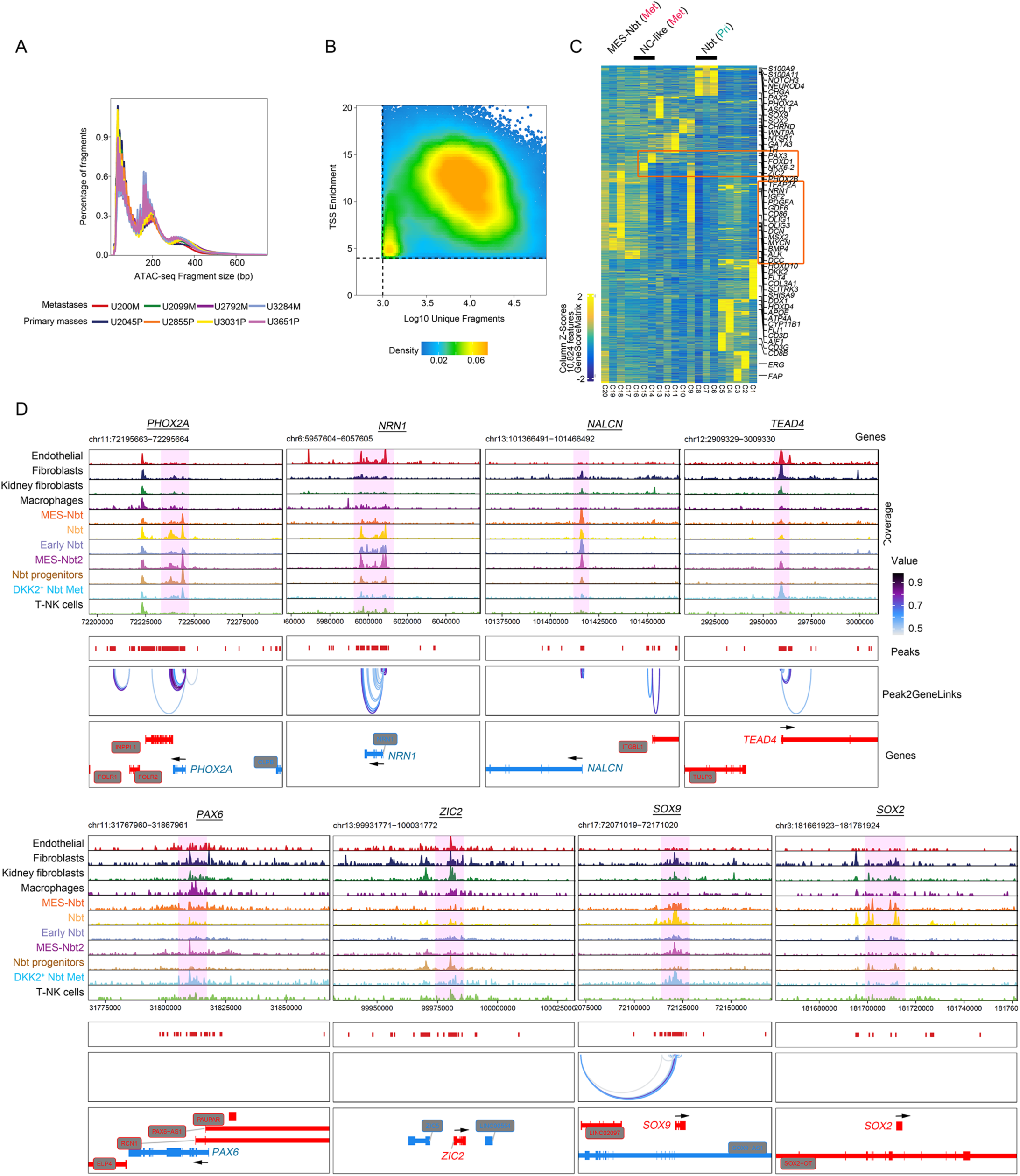
Integrative single-cell transcriptomic and epigenomic profiles reveal accessible genes in primary and metastatic NBs. a. Fragment size distributions of four primary and four metastatic NB scATAC-seq datasets. b. Transcription-start-site (TSS) enrichment scores of primary and metastatic NBs. c. Activity scores for indicated genes across cell types in primary and metastatic NBs. d. Genome browser tracks of co-accessibility at adrenergic, mesenchymal, stem cell, and metastasis marker gene loci in indicated cell types from NBs.

**Supplementary Fig. S7.**
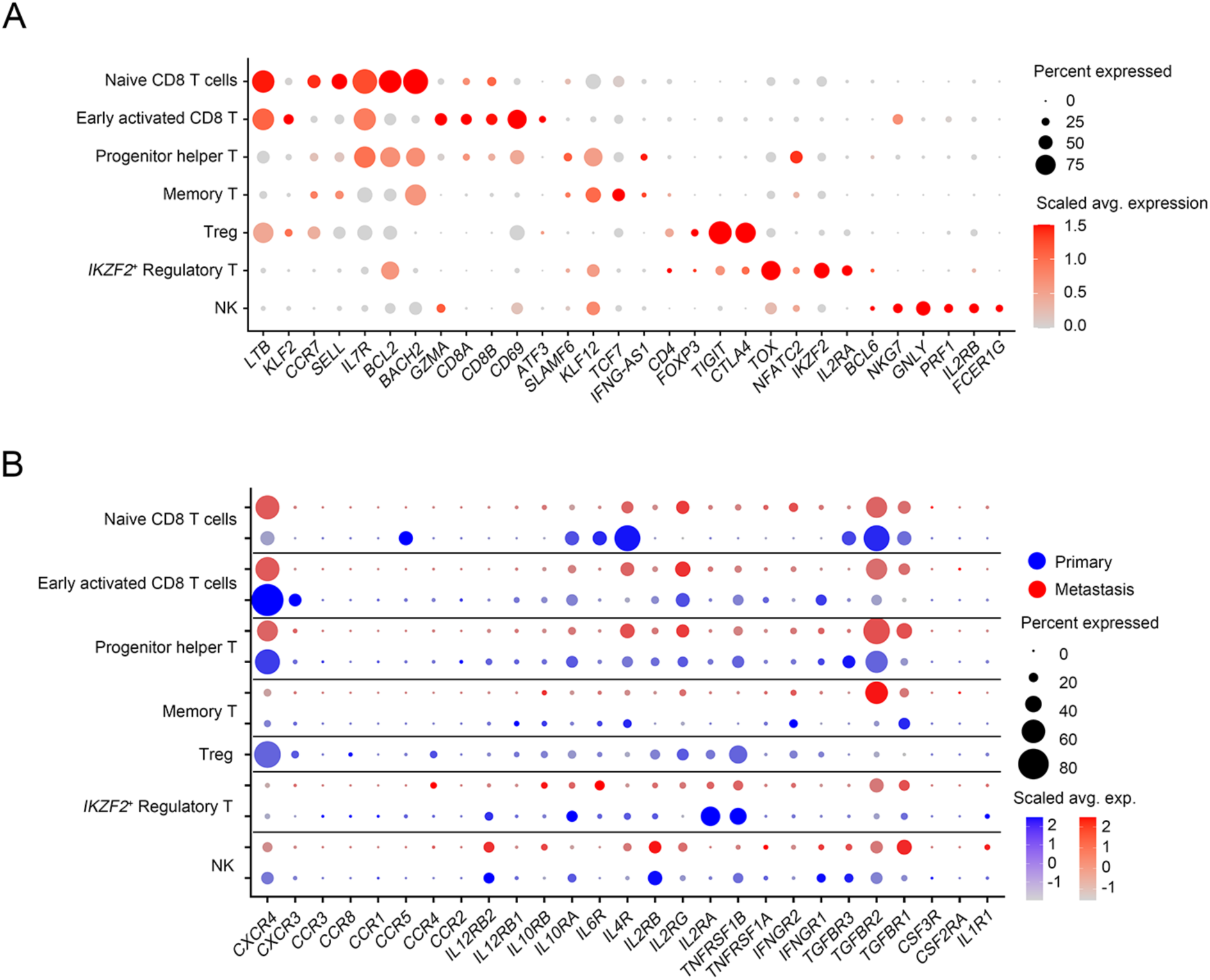
Cytokine genes are differentially expressed in lymphocytes of primary and metastatic NBs. a. Expression of indicated genes in lymphocyte clusters. b. Expression of cytokine genes primary and metastatic NBs.

**Supplementary Fig. S8.**
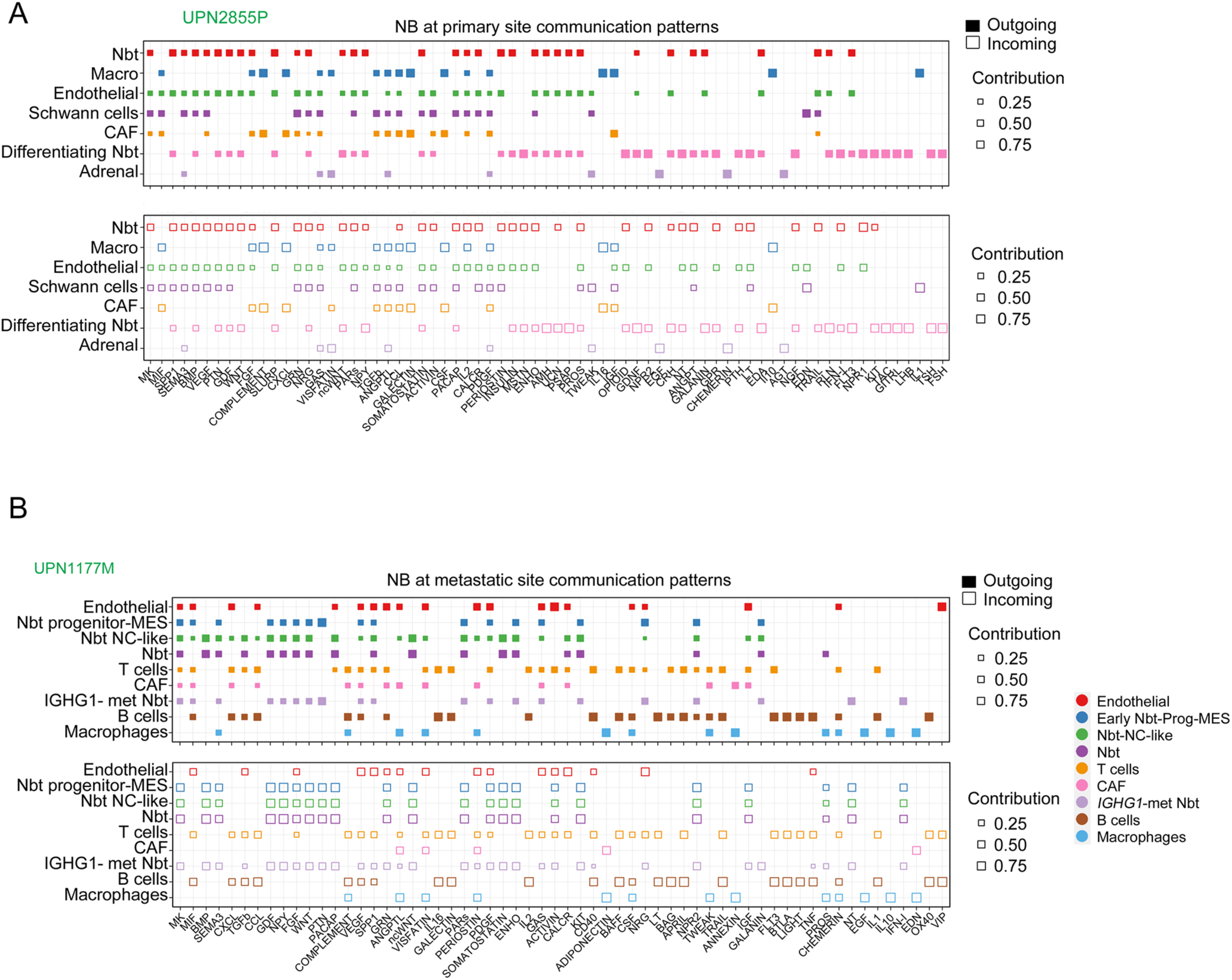
Cell-type-specific communication maps of primary NBs and lymph node metastases. a. Dot plots of outgoing (top) and incoming (bottom) signaling patterns of spatially proximal cell-cell communications between indicated cell types in primary NBs. Dot size is proportional to the contribution score computed from pattern recognition analysis using CellChat v2 ^53^. Higher contribution scores indicate more enriched signaling pathway in the corresponding cell group. b. Dot plots of outgoing (top) and incoming (bottom) signaling patterns of spatially proximal cell-cell communications between indicated cell types metastatic sites.

**Supplementary Fig. S9.**
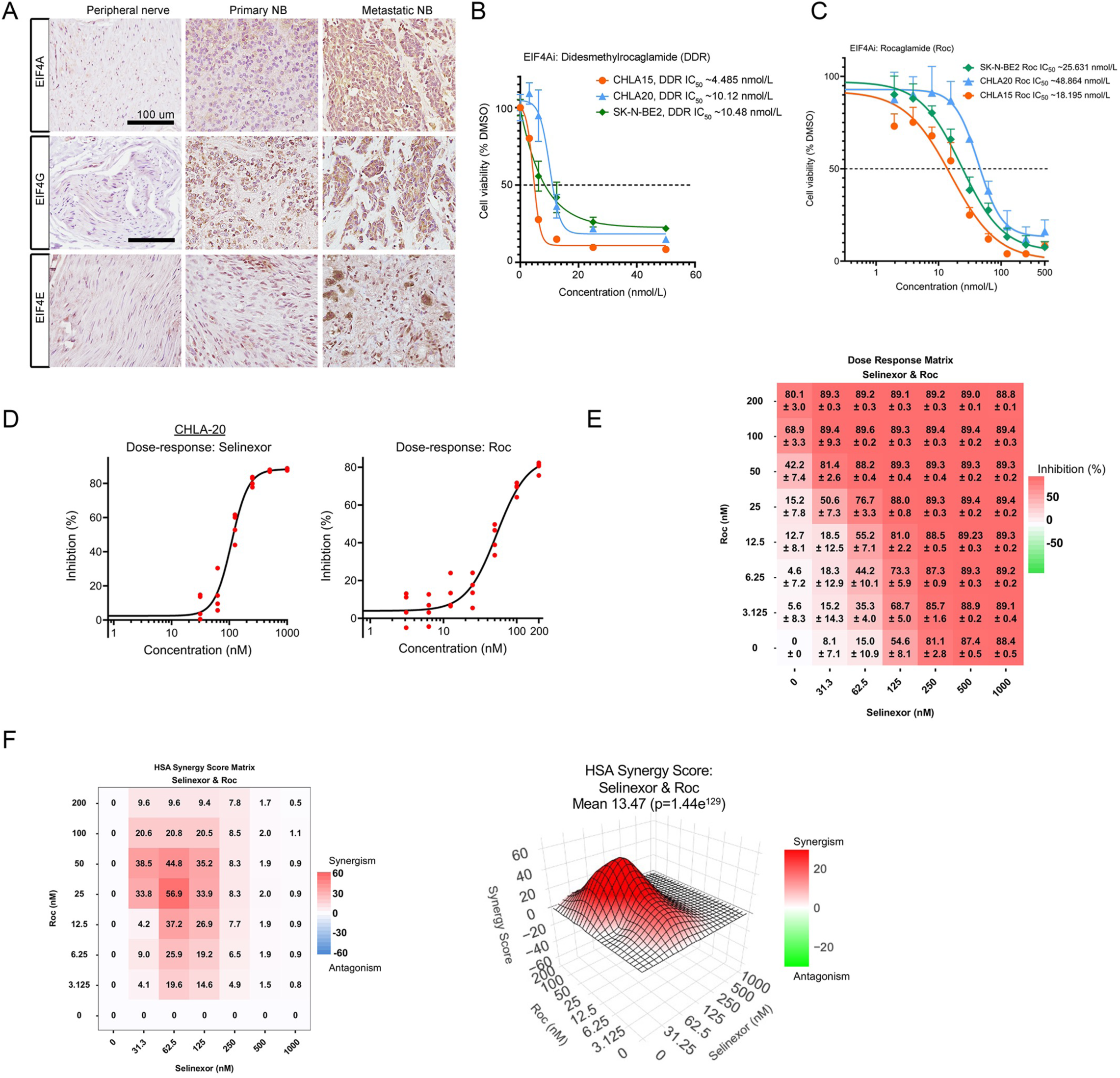
Inhibition of protein translation machinery and nuclear expert synergistically halts NB cell growth. a. Representative immunohistochemistry images of peripheral nerve, primary NB, and metastases stained for EIF4A, EIF4G, and EIF4E. Scale bars, 100 µm. b, c, Growth curves of human NB-derived cell lines CHLA15 (primary), CHLA20 (relapsed), and SK-N-BE2 (relapsed) treated with b) didesmethylrocagamide (DDR) or c) rocaglamide (Roc) (right). Cell viability was estimated as a percentage of the DMSO vehicle control. Graphs depict the means and SEMs from 2 or 3 independent experiments. Estimated IC_50_ values for each cell line are included in graph legend. d, Percent inhibition of growth of NB tumor cells as a function of Roc (left) and selinexor (right) concentrations. Red circles are means of 4 replicates. e, 2D dose response matrix showing the mean percent inhibition ± SD for each selinexor/Roc concentration combination (n=4). f. 2D matrix array (upper) and 3D contour plot (lower) of highest single agent (HSA) synergy scores calculated using SynergyFinder Plus ^92^.

## REFERENCES

1. Matthay, K.K. et al. Neuroblastoma. Nat Rev Dis Primers 2, 16078 (2016).

2. Ora, I. & Eggert, A. Progress in treatment and risk stratification of neuroblastoma: impact on future clinical and basic research. Semin Cancer Biol 21, 217–28 (2011).

3. Pinto, N.R. et al. Advances in Risk Classification and Treatment Strategies for Neuroblastoma. J Clin Oncol 33, 3008–17 (2015).

4. Bartolucci, D. et al. MYCN Impact on High-Risk Neuroblastoma: From Diagnosis and Prognosis to Targeted Treatment. Cancers (Basel) 14(2022).

5. Yanishevski, D. et al. Impact of MYCN status on response of high-risk neuroblastoma to neoadjuvant chemotherapy. J Pediatr Surg 55, 130–134 (2020).

6. Morgenstern, D.A. et al. Prognostic significance of pattern and burden of metastatic disease in patients with stage 4 neuroblastoma: A study from the International Neuroblastoma Risk Group database. Eur J Cancer 65, 1–10 (2016).

7. Yu, W. et al. Longitudinal single-cell multiomic atlas of high-risk neuroblastoma reveals chemotherapy-induced tumor microenvironment rewiring. Nat Genet 57, 1142–1154 (2025).

8. Xu, Y. et al. Single-cell MultiOmics and spatial transcriptomics demonstrate neuroblastoma developmental plasticity. Dev Cell (2025).

9. Boeva, V. et al. Heterogeneity of neuroblastoma cell identity defined by transcriptional circuitries. Nat Genet 49, 1408–1413 (2017).

10. van Groningen, T. et al. Neuroblastoma is composed of two super-enhancer-associated differentiation states. Nat Genet 49, 1261–1266 (2017).

11. Bedoya-Reina, O.C. et al. Single-nuclei transcriptomes from human adrenal gland reveal distinct cellular identities of low and high-risk neuroblastoma tumors. Nat Commun 12, 5309 (2021).

12. Dong, R. et al. Single-Cell Characterization of Malignant Phenotypes and Developmental Trajectories of Adrenal Neuroblastoma. Cancer Cell 38, 716–733 e6 (2020).

13. Fetahu, I.S. et al. Single-cell transcriptomics and epigenomics unravel the role of monocytes in neuroblastoma bone marrow metastasis. Nat Commun 14, 3620 (2023).

14. Hanemaaijer, E.S. et al. Single-cell atlas of developing murine adrenal gland reveals relation of Schwann cell precursor signature to neuroblastoma phenotype. Proc Natl Acad Sci U S A 118(2021).

15. Kameneva, P. et al. Single-cell transcriptomics of human embryos identifies multiple sympathoblast lineages with potential implications for neuroblastoma origin. Nat Genet 53, 694–706 (2021).

16. Kildisiute, G. et al. Tumor to normal single-cell mRNA comparisons reveal a pan-neuroblastoma cancer cell. Sci Adv 7(2021).

17. Ponzoni, M. et al. Recent advances in the developmental origin of neuroblastoma: an overview. J Exp Clin Cancer Res 41, 92 (2022).

18. Yuan, X. et al. Single-cell profiling of peripheral neuroblastic tumors identifies an aggressive transitional state that bridges an adrenergic-mesenchymal trajectory. Cell Rep 41, 111455 (2022).

19. Wienke, J. et al. The immune landscape of neuroblastoma: Challenges and opportunities for novel therapeutic strategies in pediatric oncology. Eur J Cancer 144, 123–150 (2021).

20. Rabadan, M.A., Usieto, S., Lavarino, C. & Marti, E. Identification of a putative transcriptome signature common to neuroblastoma and neural crest cells. Dev Neurobiol 73, 815–27 (2013).

21. Hauer, K. et al. DKK2 mediates osteolysis, invasiveness, and metastatic spread in Ewing sarcoma. Cancer Res 73, 967–77 (2013).

22. Auslander, N. et al. An integrated computational and experimental study uncovers FUT9 as a metabolic driver of colorectal cancer. Mol Syst Biol 13, 956 (2017).

23. Dong, L. et al. DIAPH3 promoted the growth, migration and metastasis of hepatocellular carcinoma cells by activating beta-catenin/TCF signaling. Mol Cell Biochem 438, 183–190 (2018).

24. Zhang, F. et al. FitDevo: accurate inference of single-cell developmental potential using sample-specific gene weight. Brief Bioinform 23(2022).

25. Street, K. et al. Slingshot: cell lineage and pseudotime inference for single-cell transcriptomics. BMC Genomics 19, 477 (2018).

26. Van den Berge, K., et al. Trajectory-based differential expression analysis for single-cell sequencing data. Nat Commun 11, 1201 (2020).

27. Yang, Q.Q. et al. Nuclear isoform of FGF13 regulates post-natal neurogenesis in the hippocampus through an epigenomic mechanism. Cell Rep 35, 109127 (2021).

28. Zhao, L., Song, W. & Chen, Y.G. Mesenchymal-epithelial interaction regulates gastrointestinal tract development in mouse embryos. Cell Rep 40, 111053 (2022).

29. El-Hashash, A.H., Al Alam, D., Turcatel, G., Bellusci, S. & Warburton, D. Eyes absent 1 (Eya1) is a critical coordinator of epithelial, mesenchymal and vascular morphogenesis in the mammalian lung. Dev Biol 350, 112–26 (2011).

30. Tirosh, I. et al. Single-cell RNA-seq supports a developmental hierarchy in human oligodendroglioma. Nature 539, 309–313 (2016).

31. Szewczyk, K. et al. Unfavorable Outcome of Neuroblastoma in Patients With 2p Gain. Front Oncol 9, 1018 (2019).

32. Depuydt, P. et al. Genomic Amplifications and Distal 6q Loss: Novel Markers for Poor Survival in High-risk Neuroblastoma Patients. J Natl Cancer Inst 110, 1084–1093 (2018).

33. Korber, V. et al. Neuroblastoma arises in early fetal development and its evolutionary duration predicts outcome. Nat Genet 55, 619–630 (2023).

34. Luttikhuis, M.E. et al. Neuroblastomas with chromosome 11q loss and single copy MYCN comprise a biologically distinct group of tumours with adverse prognosis. Br J Cancer 85, 531–7 (2001).

35. Newman, A.M. et al. Determining cell type abundance and expression from bulk tissues with digital cytometry. Nat Biotechnol 37, 773–782 (2019).

36. McCarthy, D.J., Campbell, K.R., Lun, A.T. & Wills, Q.F. Scater: pre-processing, quality control, normalization and visualization of single-cell RNA-seq data in R. Bioinformatics 33, 1179–1186 (2017).

37. Aibar, S. et al. SCENIC: single-cell regulatory network inference and clustering. Nat Methods 14, 1083–1086 (2017).

38. Martinelli, P. et al. GATA6 regulates EMT and tumour dissemination, and is a marker of response to adjuvant chemotherapy in pancreatic cancer. Gut 66, 1665–1676 (2017).

39. Sanchez-Tillo, E. et al. The EMT activator ZEB1 promotes tumor growth and determines differential response to chemotherapy in mantle cell lymphoma. Cell Death Differ 21, 247–57 (2014).

40. Li, W. et al. The FOXN3-NEAT1-SIN3A repressor complex promotes progression of hormonally responsive breast cancer. J Clin Invest 127, 3421–3440 (2017).

41. Granja, J.M. et al. ArchR is a scalable software package for integrative single-cell chromatin accessibility analysis. Nat Genet 53, 403–411 (2021).

42. Xue, J. et al. Transcriptome-based network analysis reveals a spectrum model of human macrophage activation. Immunity 40, 274–88 (2014).

43. Theruvath, J. et al. Anti-GD2 synergizes with CD47 blockade to mediate tumor eradication. Nat Med 28, 333–344 (2022).

44. Verhoeven, B.M. et al. The immune cell atlas of human neuroblastoma. Cell Rep Med 3, 100657 (2022).

45. Xia, A., Zhang, Y., Xu, J., Yin, T. & Lu, X.J. T Cell Dysfunction in Cancer Immunity and Immunotherapy. Front Immunol 10, 1719 (2019).

46. Cibrian, D. & Sanchez-Madrid, F. CD69: from activation marker to metabolic gatekeeper. Eur J Immunol 47, 946–953 (2017).

47. Khan, O. et al. TOX transcriptionally and epigenetically programs CD8(+) T cell exhaustion. Nature 571, 211–218 (2019).

48. Seo, H. et al. TOX and TOX2 transcription factors cooperate with NR4A transcription factors to impose CD8(+) T cell exhaustion. Proc Natl Acad Sci U S A 116, 12410–12415 (2019).

49. Tille, L. et al. Activation of the transcription factor NFAT5 in the tumor microenvironment enforces CD8(+) T cell exhaustion. Nat Immunol 24, 1645–1653 (2023).

50. Gabriel, S.S. et al. Transforming growth factor-beta-regulated mTOR activity preserves cellular metabolism to maintain long-term T cell responses in chronic infection. Immunity 54, 1698–1714 e5 (2021).

51. Piwecka, M., Rajewsky, N. & Rybak-Wolf, A. Single-cell and spatial transcriptomics: deciphering brain complexity in health and disease. Nat Rev Neurol 19, 346–362 (2023).

52. Joshi, S. Targeting the Tumor Microenvironment in Neuroblastoma: Recent Advances and Future Directions. Cancers (Basel) 12(2020).

53. Jin, S., Plikus, M.V. & Nie, Q. CellChat for systematic analysis of cell-cell communication from single-cell transcriptomics. Nat Protoc (2024).

54. Kishida, S. & Kadomatsu, K. Involvement of midkine in neuroblastoma tumourigenesis. Br J Pharmacol 171, 896–904 (2014).

55. Filippou, P.S., Karagiannis, G.S. & Constantinidou, A. Midkine (MDK) growth factor: a key player in cancer progression and a promising therapeutic target. Oncogene 39, 2040–2054 (2020).

56. Xiang, T., Cheng, N., Huang, B., Zhang, X. & Zeng, P. Important oncogenic and immunogenic roles of SPP1 and CSF1 in hepatocellular carcinoma. Med Oncol 40, 158 (2023).

57. Zhao, J. et al. Current insights into the expression and functions of tumor-derived immunoglobulins. Cell Death Discov 7, 148 (2021).

58. Wang, X.H. et al. IGF1R facilitates epithelial-mesenchymal transition and cancer stem cell properties in neuroblastoma via the STAT3/AKT axis. Cancer Manag Res 11, 5459–5472 (2019).

59. Silvera, D., Formenti, S.C. & Schneider, R.J. Translational control in cancer. Nat Rev Cancer 10, 254–66 (2010).

60. Bhat, M. et al. Targeting the translation machinery in cancer. Nat Rev Drug Discov 14, 261–78 (2015).

61. Kocak, H. et al. Hox-C9 activates the intrinsic pathway of apoptosis and is associated with spontaneous regression in neuroblastoma. Cell Death Dis 4, e586 (2013).

62. Chang, L.S. et al. Targeting Protein Translation by Rocaglamide and Didesmethylrocaglamide to Treat MPNST and Other Sarcomas. Mol Cancer Ther 19, 731–741 (2020).

63. Azmi, A.S., Uddin, M.H. & Mohammad, R.M. The nuclear export protein XPO1 - from biology to targeted therapy. Nat Rev Clin Oncol 18, 152–169 (2021).

64. Li, S. et al. Dual targeting of protein translation and nuclear protein export results in enhanced antimyeloma effects. Blood Adv 7, 2926–2937 (2023).

65. Lucas, J.T., Jr., et al. Risk factors associated with metastatic site failure in patients with high-risk neuroblastoma. Clin Transl Radiat Oncol 34, 42–50 (2022).

66. DuBois, S.G., Macy, M.E. & Henderson, T.O. High-Risk and Relapsed Neuroblastoma: Toward More Cures and Better Outcomes. Am Soc Clin Oncol Educ Book 42, 1–13 (2022).

67. Clere, N., Renault, S. & Corre, I. Endothelial-to-Mesenchymal Transition in Cancer. Front Cell Dev Biol 8, 747 (2020).

68. Mabe, N.W. et al. Transition to a mesenchymal state in neuroblastoma confers resistance to anti-GD2 antibody via reduced expression of ST8SIA1. Nat Cancer 3, 976–993 (2022).

69. Mei, S., et al. Single-cell analyses of metastatic bone marrow in human neuroblastoma reveals microenvironmental remodeling and metastatic signature. JCI Insight 9(2024).

70. Wang, H., Yung, M.M.H., Ngan, H.Y.S., Chan, K.K.L. & Chan, D.W. The Impact of the Tumor Microenvironment on Macrophage Polarization in Cancer Metastatic Progression. Int J Mol Sci 22(2021).

71. Qian, B.Z. & Pollard, J.W. Macrophage diversity enhances tumor progression and metastasis. Cell 141, 39–51 (2010).

72. Bellomo, C., Caja, L. & Moustakas, A. Transforming growth factor beta as regulator of cancer stemness and metastasis. Br J Cancer 115, 761–9 (2016).

73. Batlle, E. & Massague, J. Transforming Growth Factor-beta Signaling in Immunity and Cancer. Immunity 50, 924–940 (2019).

74. Ma, R., Sun, J.H. & Wang, Y.Y. The role of transforming growth factor-beta (TGF-beta) in the formation of exhausted CD8 + T cells. Clin Exp Med 24, 128 (2024).

75. Ali, M.U., Ur Rahman, M.S., Jia, Z. & Jiang, C. Eukaryotic translation initiation factors and cancer. Tumour Biol 39, 1010428317709805 (2017).

76. Smith, R.C.L. et al. Translation initiation in cancer at a glance. J Cell Sci 134(2021).

77. Pelletier, J., Graff, J., Ruggero, D. & Sonenberg, N. Targeting the eIF4F translation initiation complex: a critical nexus for cancer development. Cancer Res 75, 250–63 (2015).

78. Osborne, M.J. & Borden, K.L. The eukaryotic translation initiation factor eIF4E in the nucleus: taking the road less traveled. Immunol Rev 263, 210–23 (2015).

79. Azizian, N.G. & Li, Y. XPO1-dependent nuclear export as a target for cancer therapy. J Hematol Oncol 13, 61 (2020).

80. Patel, A.G. et al. A spatial cell atlas of neuroblastoma reveals developmental, epigenetic and spatial axis of tumor heterogeneity. bioRxiv (2024).

81. Chapple, R.H. et al. An integrated single-cell RNA-seq map of human neuroblastoma tumors and preclinical models uncovers divergent mesenchymal-like gene expression programs. Genome Biol 25, 161 (2024).

82. Hao, Y. et al. Integrated analysis of multimodal single-cell data. Cell 184, 3573–3587 e29 (2021).

83. Chen, J., Bardes, E.E., Aronow, B.J. & Jegga, A.G. ToppGene Suite for gene list enrichment analysis and candidate gene prioritization. Nucleic Acids Res 37, W305–11 (2009).

84. Stuart, T. et al. Comprehensive Integration of Single-Cell Data. Cell 177, 1888–1902 e21 (2019).

85. Emig, D. et al. AltAnalyze and DomainGraph: analyzing and visualizing exon expression data. Nucleic Acids Res 38, W755–62 (2010).

86. Venteicher, A.S. et al. Decoupling genetics, lineages, and microenvironment in IDH-mutant gliomas by single-cell RNA-seq. Science 355(2017).

87. Saelens, W., Cannoodt, R., Todorov, H. & Saeys, Y. A comparison of single-cell trajectory inference methods. Nat Biotechnol 37, 547–554 (2019).

88. Gulati, G.S. et al. Single-cell transcriptional diversity is a hallmark of developmental potential. Science 367, 405–411 (2020).

89. Hao, Y. et al. Dictionary learning for integrative, multimodal and scalable single-cell analysis. Nat Biotechnol 42, 293–304 (2024).

90. Subramanian, A. et al. Gene set enrichment analysis: a knowledge-based approach for interpreting genome-wide expression profiles. Proc Natl Acad Sci U S A 102, 15545–50 (2005).

91. Weiss, W.A., Aldape, K., Mohapatra, G., Feuerstein, B.G. & Bishop, J.M. Targeted expression of MYCN causes neuroblastoma in transgenic mice. EMBO J 16, 2985–95 (1997).

92. Zheng, S. et al. SynergyFinder Plus: Toward Better Interpretation and Annotation of Drug Combination Screening Datasets. Genomics Proteomics Bioinformatics 20, 587–596 (2022).

93. Lai, J., Tang, J., Li, T., Zhang, A. & Mao, L. Evaluating the relative importance of predictors in Generalized Additive Models using the gam.hp R package. Plant Divers 46, 542–546 (2024).

